# Astrocyte redox imbalance underlies prelimbic neuronal hypoactivity and affective behavior in epilepsy

**DOI:** 10.1101/796474

**Authors:** Travis E. Faust, Atsushi Saito, Shoichi Ishikawa, Kun Yang, Wendy Xin, Ho Namkung, Amit Agarwal, Adriana Ramos, Brian J. Lee, Lindsay Hayes, Rupali Srivastava, Sunday Adelakun, Sneha Saha, Trexy Palen, Tyler Cash-Padgett, Daniel J. Wood, Elisa Carloni, Ryosuke Yusa, Hanna Jaaro-Peled, Jed W. Fahey, Dwight E. Bergles, Koko Ishizuka, Akira Sawa

## Abstract

A fundamental but unanswered question in neuropsychiatry is whether the psychiatric symptoms of epilepsy are caused by the same or a separate pathophysiology as seizures. To address this question, we investigated a monogenic form of epilepsy (pyridoxine-dependent epilepsy) caused by *aldehyde dehydrogenase 7a1* (*ALDH7A1*) mutations. ALDH7A1 global knockout mice exhibited both seizure-associated and affective behavioral phenotypes. However, seizure phenotypes were caused by ALDH7A1 deletion in hepatocytes whereas affective behaviors were caused by ALDH7A1 deletion in astrocytes. Deletion in astrocytes disrupted astrocyte redox homeostasis, impairing regulation of extracellular ion concentrations and reducing neuronal activity in the prelimbic cortex. Sulforaphane, which activates the NRF2 antioxidant pathway, restored prelimbic neuronal activity and rescued affective behaviors in ALDH7A1 knockout mice, but did not prevent seizures. These studies implicate astrocyte redox homeostasis and prelimbic hypoactivity in the psychiatric manifestations of a congenital form of epilepsy, which are mechanistically and therapeutically separate from the seizure pathophysiology.

**Teaser:** Antioxidant imbalance in astrocytes reduces neuronal activity, causing psychiatric symptoms in epilepsy distinct from seizures.

## Introduction

Patients diagnosed with epilepsy not only suffer from seizures, but also from psychiatric and cognitive symptoms in the periods between seizures (*1, 2*). A fundamental but unanswered question is whether psychiatric and cognitive symptoms in epilepsy stem from the same pathophysiology that causes seizures or a separate pathophysiology (*3*). Epidemiological and genetic studies suggest that etiological factors are at least in part shared between epilepsy and major psychiatric disorders (*4-6*) but there is a lack of neurobiological studies on psychiatric symptoms in epilepsy to address this question. Resolving this mechanistic ambiguity will provide guidance for future treatment of psychiatric symptoms associated with epilepsy and further inform treatment of psychiatric disorders in general.

Many sporadic neuropsychiatric disorders such as epilepsy are clinically defined, which results in diagnoses that are etiologically and pathophysiologically heterogeneous. Accordingly, scientists have increasingly turned to rare genetic disorders in which the biological mechanisms can be logically studied from a single genetic etiology to clinical manifestations, which in turn provides invaluable insight to sporadic conditions (*7*). One such case is pyridoxine-dependent epilepsy (PDE) which is caused in humans by loss-of-function mutations in the *Aldehyde Dehydrogenase 7A1* (*ALDH7A1*) gene (*8*). PDE is a good model for understanding the pathophysiology of psychiatric symptoms in epilepsy since seizures in PDE are responsive to high doses of pyridoxine (also known as vitamin B6), yet psychiatric symptoms are non-responsive to pyridoxine and persist into adulthood even after seizures are controlled (*9, 10*). Furthermore, the astrocyte-specific expression pattern of *ALDH7A1* in the mature brain (*11*) facilitates molecular dissection of its cellular mechanism using genetic tools in animal models.

Astrocytes are principally responsible for homeostatic functions to support other brain cells, particularly neurons (*12*). In addition to metabolic functions, astrocytes influence neuronal activity through regulation of synaptogenesis, synaptic strength, neuronal excitability, and extracellular levels of ions and neurotransmitters (*13, 14*). Aldehyde dehydrogenase (ALDH) family genes have enriched expression in astrocytes (*11*), but the biological consequences of this restricted ALDH expression pattern remain unexplored. Enzymatically, ALDHs oxidize a wide range of endogenous and exogenous aldehydes into carboxylic acids via NAD(P)^+^-dependent reactions (*15*) with overlapping substrate specificities (*16*). As producers of NAD(P)H, ALDHs affect cellular redox homeostasis (*17-20*) which in turn influences other redox-dependent cellular activities. In many tissues and organisms, efficient aldehyde degradation by ALDHs is needed to limit toxic, electrophilic reactions between aldehydes and cellular macromolecules (*21-23*), and protect against massive oxidative stress (*24, 25*). The brain’s exposure to aldehydes may be especially high due to its high concentration of lipids and large energy demands (*26*). Although initial efforts of characterizing basic function of ALDH7A1 through loss-of-function models are just emerging (*27-30*), it remains elusive how ALDH7A1 influences astrocyte-neuron interactions and behavior.

The major psychiatry symptoms in patients with epilepsy include mood-associated manifestations, which affect more than one third of the patients (*1, 31*) including those with PDE (*32*). The current diagnosis of mood disorders includes a heterogeneous set of disease conditions in which the involvement of oxidative stress and inflammatory processes (*33, 34*), alterations of neuronal connectivity (*35*), and dysfunction of astrocytes (*36*) have all been reported. Several assays have been developed to assess mood-associated or affective behavior in rodents (*37, 38*), which have proved useful for drug discovery (*39-41*) and investigating the underlying neural circuitry of mood disorders. In one well-described circuit, rodent affective behavior is reversibly controlled by the activity of pyramidal neurons in layer 5 (L5) of the prelimbic cortex (*42*), making this a useful circuit to connect behavioral symptoms with cellular/molecular mechanisms underlying mood disorders.

In the present study, we generated a mouse model of PDE with constitutive global depletion of ALDH7A1 (KO^Global^) that exhibited increased seizure susceptibility and deficits in affective behavior. Based on the expression pattern of ALDH7A1, we also generated astrocyte-specific (cKO^Astro^) and liver-specific (cKO^Liver^) ALDH7A1 conditional knockout mice which each showed only a partial phenotype: cKO^Astro^ mice had deficits in affective behavior without any increase in seizure susceptibility whereas cKO^Liver^ mice had increased seizure susceptibility without deficits in affective behavior. To understand how astrocytes disrupt affective behavior, but not seizure phenotypes in mice, we performed RNA sequencing and identified changes in astrocyte redox homeostasis in ALDH7A1-deficient astrocytes, which we further validated by biochemistry and imaging approaches. Treating mice with dietary sulforaphane (SFN), which augmented NRF2 signaling in astrocytes, ameliorated affective behavior in cKO^Astro^ and KO^Global^ mice, supporting the significance of astrocyte redox homeostasis in affective behavior. Both dietary SFN and astrocyte-specific activation of the NRF2 pathway also ameliorated the reductions in L5 pyramidal neuron activity observed in the prelimbic cortex in cKO^Astro^ and KO^Global^ mice, which may contribute to these affective behaviors. Pharmacological, electrophysiological, and additional transcriptomic experiments suggest that astrocyte redox imbalance causes this neuronal hypoactivity by impairing astrocyte regulation of extracellular ion concentrations. Together, these data indicate an astrocyte-driven mechanism of psychiatric phenotypes in PDE that is independent of seizures. These findings may further apply to more general cases of major depressive disorder in which both astrocyte dysfunction and redox imbalance can occur (*34, 36*).

## Results

### Astrocyte ALDH7A1 contributes to affective behavior but not seizure susceptibility

We first generated mice with constitutive global deletion of *Aldh7a1* in all tissues (KO^Global^) to serve as a mouse model of PDE (**Fig 1A-B**). We validated the deletion of *Aldh7a1* in KO^Global^ mice by polymerase chain reaction (PCR) (**Fig. S1A**), and loss of ALDH7A1 protein expression by immunohistochemistry (**Fig. 1C-D, S1B**) and Western blotting (**Fig. 1E**). KO^Global^ mice were viable, maintained a normal lifespan at least into adulthood, and exhibited no differences in weight (**Fig. S1C**) or locomotion (**Fig. S1D**). Mimicking the epileptic phenotype observed in patients with PDE, KO^Global^ mice exhibited increased sensitivity to the chemoconvulsant pentylenetetrazol (PTZ) (**Fig. 1F**) which was rescued by dietary treatment with pyridoxine (**Fig S1E**), the compound used to ameliorate seizures in patients with PDE. KO^Global^ mice also displayed reduced responses in two rodent tests of affective behavior: the forced swim test (FST) (**Fig. 1G**) and the sucrose splash test (**Fig. 1H**). Thus, KO^Global^ mice model key aspects of PDE, including both seizure susceptibility and affective behavior.

**Fig. 1.**
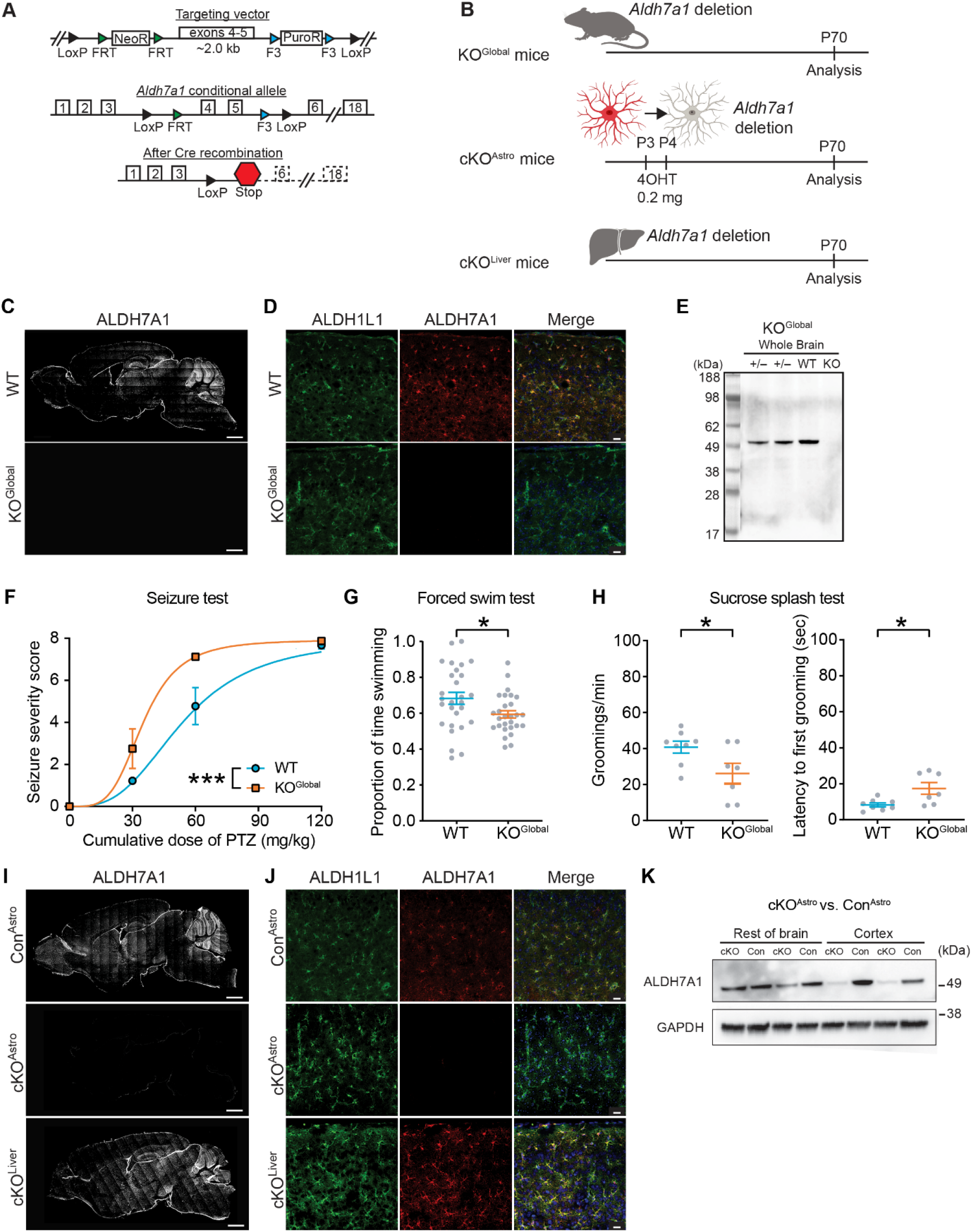
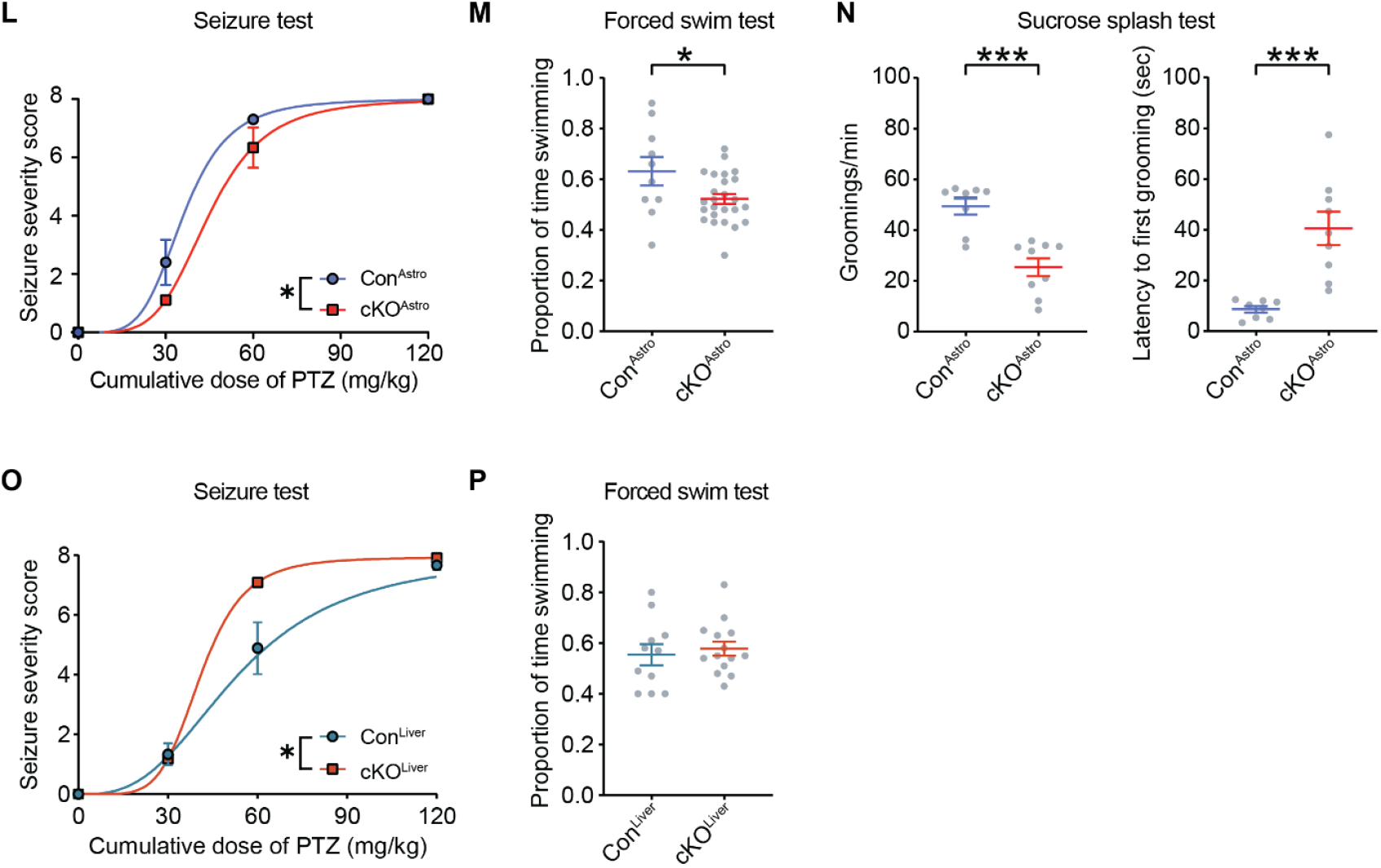
Astrocyte ALDH7A1 contributes to affective behavior but not the seizure susceptibility caused by ALDH7A1 dysfunction in the liver. (A) Schematic of targeting vector for generation of floxed *Aldh7a1* mice and floxed *Aldh7a1* allele before and after Cre recombination. (B) Schematic of strategies to generate KO^Global^, cKO^Astro^, and cKO^Liver^ mice. (C-D) Representative images of ALDH7A1 immunofluorescence in KO^Global^ and wild-type (WT) mice (C) in sagittal brain sections, scale bar 1000 µm, and (D) in the cortex, co-labeled with astrocyte marker ALDH1L1, scale bars 20 µm. (E) Western blot of ALDH7A1 in brain tissue from KO^Global^ (KO), heterozygous (+/-), and WT mice. (F) Quantification of seizure severity in KO^Global^ and WT mice in response to repeated doses of pentylenetetrazol (PTZ) during seizure threshold test. Lines represent fitted dose-response curves (Extra sum-of-squares F test on dose-response EC_50_: *n* = 8-9 mice; \*\*\**P* < 0.001). (G) Quantification of the proportion of time swimming by KO^Global^ and WT mice during the forced swim test (Student’s t-test: *n* = 28 mice; \**P* < 0.05). (H) Quantifications of frequency of grooming events (left) and latency to first grooming (right) by KO^Global^ and WT mice during the sucrose splash test (Student’s t-test: *n* = 7-8 mice; \**P* < 0.05). (I-J) Representative images of ALDH7A1 immunofluorescence in cKO^Astro^, Con^Astro^, and cKO^Liver^ mice (I) in sagittal brain sections, scale bar 1000 µm, and (J) in the cortex, co-labeled with astrocyte marker ALDH1L1, scale bars 20 µm. (K) Western blot of ALDH7A1 and GAPDH (loading control) in tissue homogenates from the cerebral cortex and the rest of the brain of cKO^Astro^ and Con^Astro^ mice. (L) Quantification of seizure severity in cKO^Astro^ and Con^Astro^ mice in response to repeated doses of PTZ during seizure threshold test. Lines represent fitted dose-response curves (Extra sum-of-squares F test on dose-response EC_50_: *n* = 9-10 mice; \**P* < 0.05). (M) Quantification of the proportion of time swimming by cKO^Astro^ and Con^Astro^ mice during the forced swim test (Student’s t-test: *n* = 10-25 mice; \**P* < 0.05). (N) Quantifications of frequency of grooming events (left) and latency to first grooming (right) by cKO^Astro^ and Con^Astro^ mice during the sucrose splash test (Student’s t-test: *n* = 8-9 mice; \*\*\**P* < 0.001). (O) Quantification of seizure severity in cKO^Liver^ and Con^Liver^ mice in response to repeated doses of pentylenetetrazol (PTZ) during seizure threshold test. Lines represent fitted dose-response curves (Extra sum-of-squares F test on dose-response EC_50_: *n* = 9-11 mice; \**P* < 0.05). (P) Quantification of proportion of time swimming by cKO^Liver^ and Con^Liver^ mice during the forced swim test (Student’s t-test: *n* = 11-14 mice; *P* > 0.05). All data represent mean ± S.E.M.

We next examined the cellular expression pattern of ALDH7A1, using KO^Global^ mice as the negative control. ALDH7A1 was expressed in astrocytes throughout the brain, but not in other major brain cell types (**Fig. S2A-J**). ALDH7A1 expression was also higher in adult mice compared to weanlings (**Fig. S2K-L)**, but did not differ between males and females (**Fig. S2M-N**). Outside the brain, ALDH7A1 was expressed in the kidney and liver (**Fig S1B**), but not other tissues. Based on the expression profile of ALDH7A1, we decided to generate mice with conditional deletion of *Aldh7a1* in astrocytes (cKO^Astro^ mice) using *Slc1a3*^CreER^ and mice with conditional deletion of *Aldh7a1* in the liver (cKO^Liver^ mice) using *Alb1*^*Cre*^ (**Fig 1B**). Immunohistochemical assessment (**Fig. 1I-J**) and Western blotting (**Fig. 1K**) confirmed that ALDH7A1 was efficiently depleted from astrocytes in cKO^Astro^ mice (∼90% in the cortex and 50% in the rest of the brain) and maintained in peripheral organs (**Fig. S1B**). In cKO^Liver^ mice, ALDH7A1 was efficiently depleted from the liver (**Fig. S1B)** and was maintained within the brain (**Fig. 11-J)**.

Comparing the behavior of cKO^Astro^ mice to KO^Global^ mice, we paradoxically observed a higher seizure threshold (i.e., a lower seizure susceptibility) in cKO^Astro^ mice compared to Cre-negative littermate controls (Con^Astro^) (**Fig. 1L**), opposite to the effect observed in KO^Global^ mice. To confirm this result, we also assessed seizure threshold in mice with constitutive, embryonic deletion of *Aldh7a1* in all forebrain astrocytes and neurons (*Emx1*^Cre/*+*^; *Aldh7a1*^flox/flox^ mice) (*43*) and again found a higher seizure threshold (**Fig. S1G**). These results suggested that depletion of ALDH7A1 from forebrain astrocytes does not drive the increased seizure susceptibility observed in KO^Global^ mice. However, despite the lack of seizure susceptibility, cKO^Astro^ mice still phenocopied the affective behavior of KO^Global^ mice with reduced mobility in the FST (**Fig. 1M**) and reduced grooming frequency and increased grooming latency during the sucrose splash test, **Fig. 1N**). These results suggest that astrocytes contribute to the altered affective behavior in KO^Global^ mice but not the increased seizure susceptibility.

In contrast to cKO^Astro^ mice, cKO^Liver^ mice had a lower seizure threshold (i.e. higher seizure susceptibility) compared to Con^Liver^ mice, phenocopying the result observed in KO^Global^ mice (**Fig. 1O**). However, cKO^Liver^ mice did not show any significant difference in mobility during the FST (**Fig. 1P**). These data suggest that ALDH7A1 depletion in the liver, rather than in astrocytes, drives the increase in seizure susceptibility in KO^Global^ mice and that ALDH7A1 depletion in astrocytes, but not the liver, is responsible for the deficits in affective behavior in KO^Global^ mice.

### Astrocyte ALDH7A1 regulates cellular redox homeostasis

In patients with PDE, seizures can be controlled by pyridoxine, but psychiatric symptoms remain intractable. We therefore focused on determining how ALDH7A1 depletion in astrocytes drives affective behavior. Although several functions of ALDH7A1 have been described in other tissues (*19, 44*), the function of ALDH7A1 in the brain has not been determined. To investigate how ALDH7A1 depletion leads to brain dysfunction, we first performed RNA sequencing (RNA-Seq) on brain tissue homogenate from KO^Global^ mice (**Fig. 2A**), focusing on the cortex where ALDH7A1 depletion was most robust in cKO^Astro^ mice (**Fig 1J**). As expected, *Aldh7a1* expression was significantly decreased in the brain of KO^Global^ mice, but only 3 other genes (*Hist1h1c, Prelp, Hist1h1e*) reached gene-level significance (adjusted *p* < 0.05; **Fig. 2B; Table S1**), suggesting that the brain transcriptome is largely unaltered by ALDH7A1 depletion. Nevertheless, gene set enrichment analysis (GSEA) of all detected genes identified 338 affected pathways (**Table S1**), which we clustered into 38 biological themes using EnrichmentMap (*45*) (**Fig. 2C**). Interestingly, although most PDE research has focused on the role of ALDH7A1 in lysine degradation (*8, 46*) a majority of the themes that were downregulated in KO^Global^ brains were related to cellular redox processes (e.g. NADH dehydrogenase activity, glutathione binding, cellular respiration, and lipid catabolism). To validate these findings, we performed biochemical assays to assess cellular redox status in tissue homogenate from KO^Global^ mice. Compared to wild-type (WT) littermates, KO^Global^ mice had a lower NAD^+^/NADH ratio in cortical tissue homogenates (**Fig. 2D-E**). For NADP^+^/NAPDH ratio and GSH/GSSG ratio, we further restricted our analysis to the prelimbic cortex (**Fig. 2F**), a region of the cortex known to regulate affective behavior during the FST (*42*) and found no change in the NADP^+^/NADPH ratio (**Fig. 2G**) but a lower ratio of reduced glutathione (GSH) to oxidized glutathione (GSSG) (**Fig. 2H**). These results suggest that ALDH7A1 significantly impacts redox homeostasis in the brain.

**Fig. 2.**
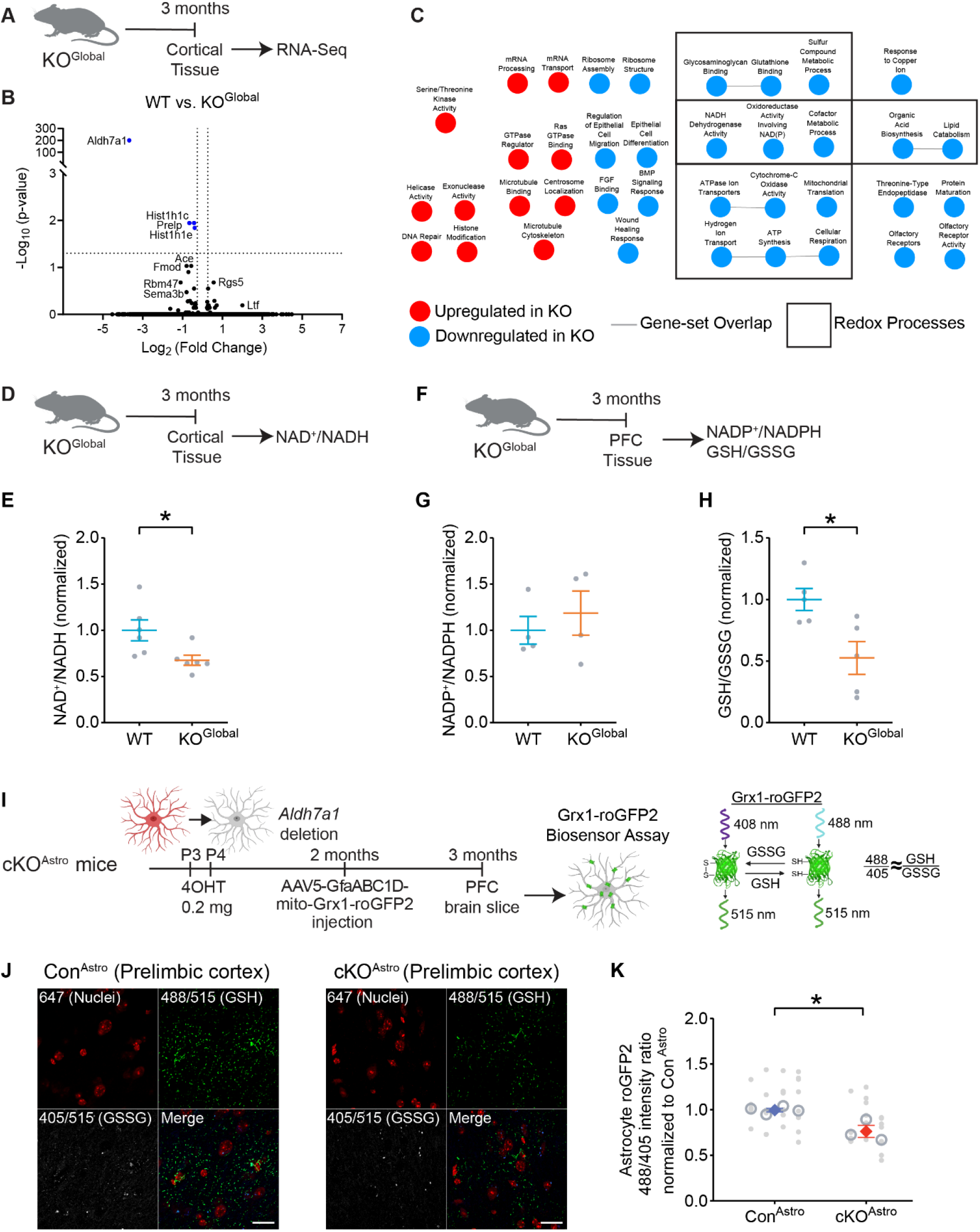
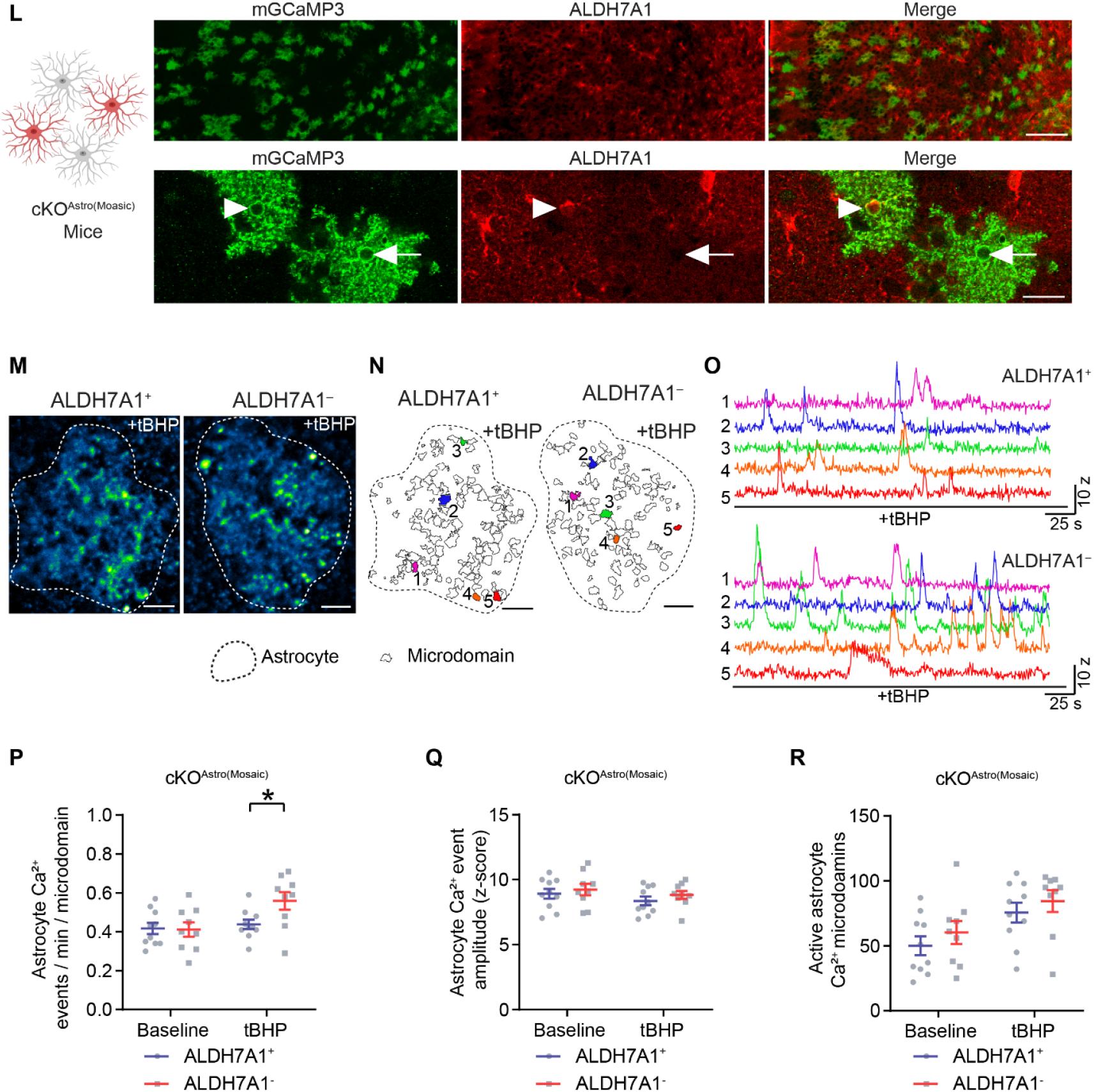
Astrocyte ALDH7A1 regulates cellular redox homeostasis. (A-B) (A) Schematic outline and (B) Volcano plot of RNA-sequencing results from KO^Global^ and wild-type (WT) mice. Dotted lines indicate cutoffs at *P* < 0.05 and log_2_ fold change > | 1.2 |. (C) EnrichmentMap summary of gene set enrichment analysis (GSEA) of KO^Global^ mice cortical tissue. Biological themes associated with redox processes are outlined. (D-E) (D) Schematic outline and (E) quantification of the NAD^+^/NADH ratio in cortical tissue homogenate from KO^Global^ and WT mice (Student’s t-test: *n* = 6 mice; \**P* < 0.05). (F-H) (F) Schematic outline and quantifications of (G) the NADP^+^/NADPH ratio and (H) the reduced (GSH) to oxidized (GSSG) glutathione ratio in tissue homogenate from the prefrontal cortex (PFC) of KO^Global^ and WT mice littermates (Student’s t-tests: NADP^+^/NAPDH: *n* = 4 mice; *P* > 0.05; GSH/GSSG: *n* = 5 mice; \**P* < 0.05). (I) Schematic of mito-Grx1-roGFP2 ratiometric biosensor assay to assess the redox ratio in astrocytes in the prelimbic cortex of cKO^Astro^ mice. (J) Representative images of 405/515 nm fluorescence (GSSG), 488/515 nm fluorescence (GSH) and TO-PRO iodide (nuclei) in cKO^Astro^ and Con^Astro^ mice expressing mito-Grx1-roGFP2 in astrocytes. Scale bars 20 μm. (K) Quantification of the mito-Grx1-roGFP2 488/405 nm fluorescence intensity ratio in cKO^Astro^ and Con^Astro^ mice. (nested t-test: *n* = 3-4 mice (open circles), 5-9 images per mouse (filled circles); \**P* < 0.05). (L) (left) Schematic ALDH7A1 depletion in cKO^Astro(Mosaic)^ mice. (Right) Representative images of ALDH7A1 and mGCaMP3 immunofluorescence in cKO^Astro(Mosaic)^ mice, scale bars: 200 µm (top), 20 μm (bottom). ALDH7A1+ (arrowhead) and ALDH7A1^−^ (arrow) astrocytes expressing mGCaMP3 are indicated. (M) Median intensity projection image (pseudocolored) of 540 frames of one ALDH7A1^+^ astrocyte (outlined; left) and one ALDH7A1^−^ astrocyte (right) after incubation with tert-butyl hydrogen peroxide (tBHP). Scale bar, 10 μm. (N) Map of microdomains recorded in one ALDH7A1^+^ astrocyte (left) and one ALDH7A1^−^ (right) after tBHP incubation. Outlines indicate cell border. Scale bar, 10 μm. (O) Intensity vs. time traces for 5 microdomains (corresponding to colors in F) depicting Ca^2+^ transients in ALDH7A1^+^ and ALDH7A1^−^ astrocytes after tBHP incubation. (P-R) Quantification of (P) the frequency of microdomain Ca^2+^ transients, (Q) Ca^2+^ transient amplitudes, and (R) the number of active microdomains in ALDH7A1^+^ and ALDH7A1^−^ astrocytes at baseline and after tBHP incubation (Repeated-measures 2-way ANOVA with Holm-Sidak post-hoc tests: *n* = 9-10 cells; \**P* < 0.05). Data in (E-H), (K), and (P-R) represent mean ± S.E.M.

Because ALDH7A1 is directly involved in degradation of reactive aldehydes and NAD^+^-dependent reactions, we hypothesized that brain redox homeostasis could be disrupted in ALDH7A1 KO^Global^ mice due to redox dysfunction in astrocytes. To selectively examine redox changes in astrocytes, we engineered an adeno-associated virus (AAV) to selectively express a ratiometric redox sensor (mito-Grx1*-*roGFP2) (*47*) in astrocytes and injected it into the prelimbic cortex of cKO^Astro^ mice (**Fig. 2I**). Compared to Con^Astro^ mice, cKO^Astro^ mice had a lower 488/405 nm ratio, indicative of a reduced GSH/GSSG ratio in astrocytes (**Fig. 2J-K**). This result matches the reduced GSH/GSSG ratio observed in tissue homogenate from KO^Global^ mice and suggests that ALDH7A1 depletion in astrocytes disrupts astrocyte redox homeostasis.

We then investigated the cell autonomous consequences of impaired redox homeostasis on redox-related functions in astrocytes. First, we looked for evidence of oxidative stress or mitochondrial dysfunction in astrocytes. Using CellROX, a cell-permeable dye that fluoresces upon oxidation, we did not detect significant increases in reactive oxygen species (ROS) in astrocytes by flow cytometry (**Fig. S3A-C**). We also performed Seahorse analysis of cellular respiration in primary astrocytes and found no changes in cellular respiration at baseline or under mitochondrial stress (**Fig. S3D**). Taken together, these results suggested that global ALDH7A1 depletion did not elicit robust oxidative stress in astrocytes at the levels frequently associated with cell and tissue damage.

Next, we assessed the cell autonomous effects of ALDH7A1 depletion on astrocyte redox homeostasis *in vivo*. For these experiments, we used a “mosaic deletion” strategy (*48*) to generate mice with *Aldh7a1*-deleted and -undeleted astrocytes intermixed within the same animal [*Aldh7a1*^*flox/flox*^; BAC*-Slc1a3*^*CreER*^; *R26-LSL-mGCaMP3*^*flox/+*^ mice (cKO^Astro(Mosaic)^)]. By using a less efficient strain of *Slc1a3*^CreER^ (Tg(Slc1a3-cre/ERT)1Nat/J), we were able to delete *Aldh7a1* in ∼50% of cortical astrocytes and sparsely label ALDH7A1-immunopositive (ALDH7A1^+^) and - immunonegative (ALDH7A1^−^) astrocytes with membrane-bound GCaMP3, an effective readout of ROS-sensitive Ca^2+^ signaling events in astrocyte processes (*49*) (**Fig. 2L and S3E**). ALDH7A1^+^ and ALDH7A1^−^ astrocytes had a similar expression of an astrocyte reactivity marker, glial fibrillary acidic protein (GFAP), and an oxidative stress marker, 8-oxo-dG (**Fig. S3F-G**), which suggested no gross changes in cellular reactivity or robust oxidative stress occur in astrocytes *in vivo*. We then assessed astrocyte intracellular Ca^2+^ signaling in acute cortical brain slices from the prelimbic cortex (**Fig. 2M-O**). At baseline, neither the frequency nor amplitude of Ca^2+^ transients (**Fig. 2P-Q**), nor the number of active microdomains (**Fig. 2R**), were different between ALDH7A1^+^ and ALDH7A1^−^ cells, suggesting that ALDH7A1^−^ cells did not have altered Ca^2+^ signaling under normative conditions. However, following brief, low-dose exposure to tert-butyl hydrogen peroxide, a compound that stimulates ROS production (*50*), the frequency of Ca^2+^ transients was significantly higher in ALDH7A1^−^ cells than ALDH7A1^+^ cells (**Fig. 2P**). These results indicated that ALDH7A1-deficient astrocytes had an increased sensitivity to redox imbalance. Taken together, these results implied a role of ALDH7A1 in astrocyte redox homeostasis at both molecular expression (**Fig. 2A-K; Table S1**) and functional (**Fig. 2L-R**) levels, unaccompanied by robust oxidative stress or cell damage (**Fig. S3A-G**).

### SFN ameliorates affective behavior, but not seizure susceptibility, in ALDH7A1 knockout mice

To determine if astrocyte redox imbalance is a significant contributor to brain and behavioral phenotypes in ALDH7A1 knockout mice, we tested the impact of treating mice with sulforaphane (SFN), a brain-penetrant antioxidant derived from broccoli sprouts. SFN helps cells maintain redox homeostasis by inducing expression of cytoprotective, catalytic enzymes via the NRF2 pathway (*51, 52*). To administer SFN, we developed a dietary formulation that remained stable and bioactive in rodent chow (**Fig. S4A**). At baseline, cKO^Astro^ mice had reduced NRF2 protein levels in astrocytes, but 1-week dietary administration of SFN was sufficient to normalize NRF2 levels (**Fig S4B-E**), suggesting that dietary SFN can successfully increase antioxidant protection in ALDH7A1-deficient astrocytes.

We then investigated the impact of long-term dietary SFN administration on the behavior of cKO^Astro^ and KO^Global^ mice (**Fig 3A-B**). Importantly, this long-term administration of SFN from 3 weeks of age until adulthood was well tolerated and did not affect animal body weight (**Fig. S5A-B**). In both cKO^Astro^ and KO^Global^ mice, SFN treatment normalized the affective behavioral deficits. When treated with SFN, neither cKO^Astro^ mice nor KO^Global^ mice exhibited abnormal FST behavior (**Fig. 3C-D, S5C-D**) or abnormal sucrose splash test behavior (**Fig. 3E-H)**. In contrast, SFN differentially influenced seizure threshold in cKO^Astro^ and KO^Global^ mice. The lower seizure susceptibility observed in cKO^Astro^ mice was normalized by SFN (**Fig. 3I, S5E**), but KO^Global^ mice still exhibited increased seizure susceptibility in the presence of SFN (**Fig. 3J, S5F**), indicating that SFN was ineffective for treating the seizure-associated phenotype in KO^Global^ mice. Together, these data suggest that SFN rescued the astrocyte-driven mechanisms of ALDH7A1 dysfunction, resulting in amelioration of astrocyte-driven affective behavior deficits, but not liver-driven mechanisms that increase seizure susceptibility.

**Fig. 3.**
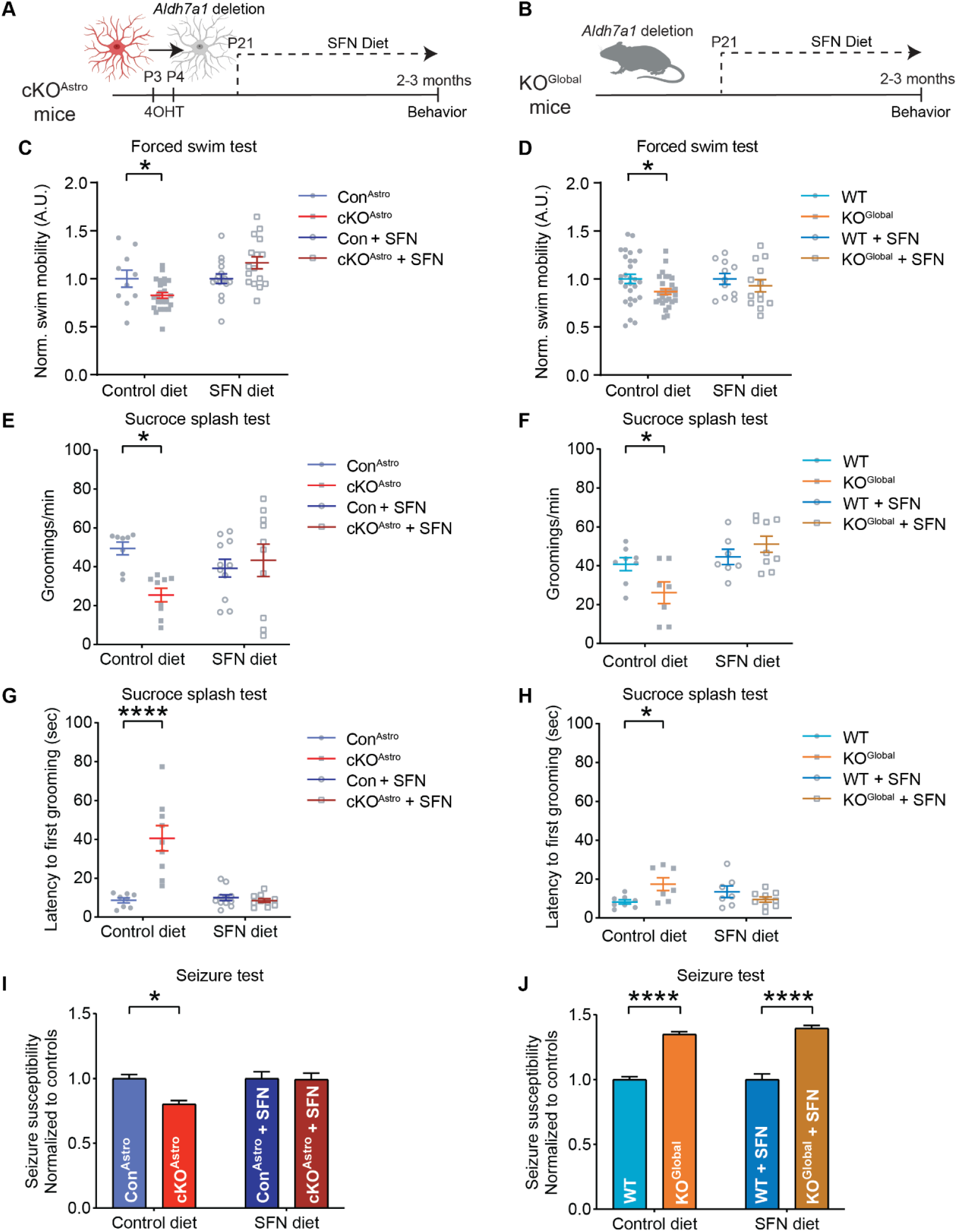
Sulforaphane ameliorates affective behavior, but not seizure susceptibility, in ALDH7A1 knockout mice. (A-B) Schematic of behavioral assessment in (A) cKO^Astro^ mice and (B) KO^Global^ mice after long-term dietary treatment with sulforaphane (SFN). (C-D) Quantifications of the proportion of time swimming by (C) cKO^Astro^ vs. Con^Astro^ mice and (D) KO^Global^ vs. WT mice on control diet or SFN diet. Data shown is from Figs. 1G, 1M, and S5C-D, normalized to respective dietary controls. (E-H) Quantifications of sucrose splash test behavior on control diet or SFN diet by cKO^Astro^ vs. Con^Astro^ mice and KO^Global^ vs. WT mice. Control diet data is from Figs. 1H, 1N. Left graphs show (E) frequency of grooming events and (G) latency to first grooming in cKO^Astro^ vs. Con^Astro^ mice (2-way ANOVA with Holm-Sidak post-hoc tests: *n* = 8-11 mice; \**P* < 0.05, \*\*\*\**P* < 0.0001). Right graphs show (F) frequency of grooming events and (H) latency to first grooming in KO^Global^ vs. WT mice (2-way ANOVA with Holm-Sidak post-hoc tests: *n* = 7-9 mice; \**P* < 0.05). (I-J) Quantification of seizure susceptibility to pentylenetetrazol (PTZ) on control diet and SFN diet by (I) cKO^Astro^ vs. Con^Astro^ and (J) KO^Global^ vs. WT mice. Data represent EC_50-1_ of PTZ dose-response curves from Figs. 1F, 1L, and S5E-F, normalized to dietary controls (2-way ANOVA with Holm-Sidak post-hoc tests. cKO^Astro^: *n* = 5-8 mice, KO^Global^: *n* = 8-12 mice; \**P* < 0.05, \*\*\*\**P* < 0.0001). All data represent mean ± S.E.M.

### SFN ameliorates astrocyte ALDH7A1-induced neuronal hypoactivity in the prelimbic cortex

Since SFN was effective in rescuing affective behaviors in both KO^Global^ and cKO^Astro^ mice, we next looked for the neural circuit that might connect astrocyte redox dyshomeostasis with deficits in affective behavior. Although multiple brain regions are involved in affective behavior, we prioritized the prelimbic cortex, a region known to regulate FST behavior (*42*) where we had observed both strong depletions in ALDH7A1 (**Fig. 1J**) and deficiencies in redox homeostasis (**Fig. 2F-R**). Pharmacological lesioning experiments using ibotenic acid further suggested that the prelimbic cortex is also required for affective behavior during the sucrose splash test (**Fig S6**). We therefore hypothesized that the prelimbic cortex may contribute to the affective behavior deficits in KO^Global^ and cKO^Astro^ mice.

The specific neurons in the prelimbic cortex that control FST swimming activity are the layer 5 (L5) pyramidal neurons that project to the brainstem (*42*), with reductions in neuronal activity causing reduced swim mobility. To examine the activity of L5 pyramidal neurons in cKO^Astro^ and KO^Global^, we used AAV9-CaMKIIa-GCaMP6f to drive L5 pyramidal neuron expression of the fluorescent Ca^2+^ indicator GCaMP6f (**Fig. 4A-B**). Notably, when expressed here in neurons, instead of in astrocytes, spikes in GCaMP fluorescence serve as indicators of neuronal activity. Therefore, to assess neuronal activity, we prepared acute brain sections from the prelimbic cortex and imaged spontaneous changes in GCaMP6f fluorescence in the soma of layer 5 pyramidal neurons. In both cKO^Astro^ mice (**Fig. 4C-D**) and KO^Global^ mice (**Fig. S7)**, there was a reduced frequency of somatic Ca^2+^ events in prelimbic L5 pyramidal neurons, suggesting that these neurons were hypoactive. We then measured Ca^2+^ event frequency in prelimbic L5 pyramidal neurons in brain slices from SFN-fed cKO^Astro^ mice. Similar to the effect of SFN diet on affective behaviors *in vivo*, the Ca^2+^ event frequency of the prelimbic L5 pyramidal neurons in cKO^Astro^ brain slices was normalized by SFN diet (**Fig. 4C-D**). Thus, across cKO^Astro^, KO^Global^, and SFN-treated cKO^Astro^ mice, the activity of L5 prelimbic pyramidal neurons correlated with affective behavior changes, suggesting that hypoactivity of L5 prelimbic pyramidal neurons underlies affective behavior changes in mice with ALDH7A1 depletion.

**Fig. 4.**
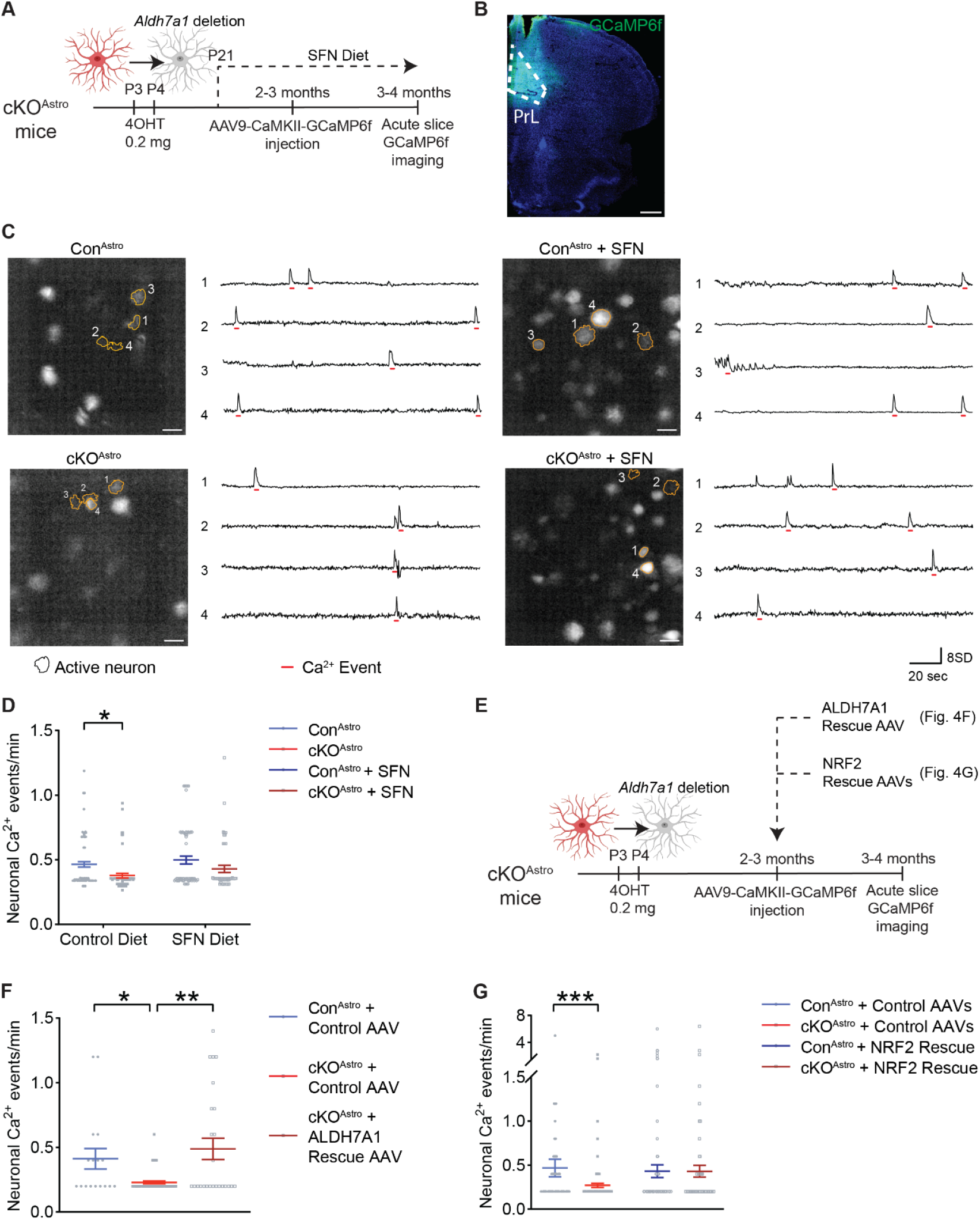
SFN ameliorates astrocyte ALDH7A1-induced neuronal hypoactivity in the prelimbic cortex. (A) Schematic of experimental strategy to assess prelimbic layer 5 (L5) pyramidal neuron activity in cKO^Astro^ mice by GCaMP6f fluorescence after long-term dietary treatment with sulforaphane (SFN). (B) Representative image of GCaMP6f fluorescence in cKO^Astro^ mice with stereotactic injection of AAV9-CaMKII-GCaMP6f in prelimbic cortex (PrL). Scale bar, 500 μm (C) Representative images (left) and traces (right) of GCaMP6f fluorescence in prelimbic L5 pyramidal neurons in cKO^Astro^ and Con^Astro^ mice on control diet and SFN diet. Numbered orange outlines indicate neurons in images used for analysis and shown in traces. Events are indicated by red underlines. Scale bars, 20 μm. (D) Quantification of prelimbic L5 pyramidal neuron Ca^2+^ event frequency in cKO^Astro^ and Con^Astro^ mice on control diet and SFN diet (Mann-Whitney with Holm-Sidak multiple comparisons test. Control diet: *n* = 63-101 neurons; SFN diet: *n* = 49-63 neurons. \**P* < 0.05). (E) Schematic overview of experimental design for (F-G). (F) Quantification of prelimbic L5 pyramidal neuron Ca^2+^ event frequency in Con^Astro^ and cKO^Astro^ mice with stereotactic injection of ALDH7A1 Rescue AAV (AAV5-GfaABC1D-ALDH7A1-mCherry) or control AAV(AAV5-GfaABC1D-mCherry). Kruskal-Wallis test with Dunn’s post-hoc tests: *n* = 17-62 neurons. *\*P* < 0.05, \*\**P* < 0.01. (G) Quantification of prelimbic L5 pyramidal neuron Ca^2+^ event frequency in Con^Astro^ and cKO^Astro^ mice with stereotactic injection of NRF2 Rescue AAVs (AAV5-GfaABC1D-Nrf2-mCherry and AAV5-GfaABC1D-*Keap1* shRNA) or control AAVs (AAV5-GfaABC1D-mCherry and AAV5-GfaABC1D-scrambled shRNA). Mann-Whitney with Holm-Sidak multiple comparisons test: control diet: *n* = 50-112 neurons; SFN diet: *n* = 105-115 neurons. \*\*\**P* < 0.001 All data represent mean ± S.E.M.

The prelimbic L5 pyramidal neuron hypoactivity is consistent with the affective behavior changes, but it is somewhat counterintuitive that neurons would be hypoactive in KO^Global^ mice that are more susceptible to seizures. To further confirm the reduced activity of prelimbic L5 pyramidal neurons and interrogate the contribution of these neurons to PTZ-induced seizures, we performed immunostaining for c-FOS, a marker of neuronal activity, at 1 h after seizures in KO^Global^ and WT mice. For consistency, we only included mice with a seizure severity score of 7 on the modified Racine scale. Compared to the dentate gyrus where ∼90% of cells were c-FOS^+^ in both WT and KO^Global^ mice, only ∼1-3% of cells were c-FOS^+^ in layer 5 of the prelimbic cortex, suggesting that the prelimbic cortex is not a significant driver of seizure activity (**Fig. S8**). In addition, the number of c-FOS^+^ cells in L5 prelimbic cortex was significantly lower in KO^Global^ mice than in WT mice (**Fig. S8F-G**), further confirming our findings that neuronal activity is reduced in the prelimbic cortex.

Having identified prelimbic L5 pyramidal neurons as an SFN-responsive neural correlate of affective behavior in ALDH7A1 knockout mice, we sought to determine whether the change in prelimbic L5 pyramidal neuron activity is due to synaptic changes or the electrical properties of the neurons. To assess synaptic inputs on prelimbic L5 pyramidal neurons, we recorded excitatory and inhibitory post-synaptic currents (ESPCs and IPSCs) from prelimbic L5 pyramidal neurons. We observed no difference in the frequency or amplitude of spontaneous EPSCs and IPSCs in cKO^Astro^ mice (**Fig. S9A-D**) and KO^Global^ mice (**Fig. S9E-H**), suggesting that prelimbic L5 pyramidal neuron hypoactivity is not due to changes in synaptic inputs. We next measured the intrinsic electrical properties of prelimbic L5 pyramidal neurons in KO^Global^ mice. Action potential shape was unaffected, as were capacitance, and membrane resistance (**Fig. S10A-C**). However, the resting membrane voltage of the neurons was decreased (**Fig. S10D**), which can lead to fewer spikes given a fixed amount of synaptic input. Indeed, we observed fewer action potentials were induced by the same current injection (**Fig. S10E**), suggesting that the prelimbic L5 pyramidal neuron hypoactivity in ALDH7A1 knockout mice was due to a reduction in neuronal excitability.

Next, we investigated how dietary SFN restores neuronal activity in ALDH7A1-deficient mice. Immunolabeling revealing increased NRF2 in astrocytes after dietary SFN (**Fig. S4C-E**), suggesting that SFN acts by restoring redox imbalance in astrocytes. To test whether restoring astrocyte function alone is sufficient to normalize neuronal activity, we generated an astrocyte-specific ALDH7A1 Rescue AAV (AAV5-GfaABC1D-ALDH7A1-mCherry) and injected it into the prelimbic cortex of adult cKO^Astro^ mice (**Fig. S11A**). The virus was expressed selectively in astrocytes (**Fig. S11B**) and restored ALDH7A1 protein levels (**Fig. S11C**). In cKO^Astro^ mice injected with control AAV, neuronal Ca^2+^ event frequency was reduced compared to Con^Astro^ mice, whereas ALDH7A1 Rescue AAV restored Ca^2+^ event frequency to control levels (**Fig. 4E-F**). Thus, neuronal hypoactivity can be reversed by correcting astrocyte ALDH7A1 deficiency, demonstrating that astrocyte-dependent regulation of neuronal activity remains reversible in adulthood.

To further test whether SFN restores neuronal activity by normalizing astrocyte redox signaling via NRF2 signaling, we used a complementary genetic strategy. NRF2 protein stability and its function as a master transcription factor are regulated by KEAP1 which targets NRF2 for ubiquitination and degradation (*53*). SFN inhibits KEAP1, thereby preventing NRF2 degradation and sustaining NRF2 signaling (*54*). To mimic the effects of SFN on NRF2 signaling, we generated two astrocyte-specific NRF2 Rescue AAVs: one to overexpress *Nrf2* (GfaABC1D-Nrf2-mCherry) and one to knock down *Keap1* (GfaABC1D-mCherry-*Keap1* shRNA) (**Fig. S11A**). Co-expression of both constructs increased NRF2 levels in astrocytes (**Fig. S11D**), confirming the validity of this approach. Similarly to the ALDH7A1 Rescue AAV, injection of NRF2 Rescue AAVs into the prelimbic cortex of cKO^Astro^ restored neuronal Ca^2+^ event frequency to control levels (**Fig 4E, G**). Together, these findings show that prelimbic neuronal hypoactivity results from astrocyte redox dysregulation caused by ALDH7A1 deficiency and can be reversibly corrected by restoring ALDH7A1 or activating the NRF2 pathway in astrocytes, including through dietary SFN.

### SFN ameliorates expression of ion channel and ion transporter gene pathways in cKO^**Astro**^ astrocytes

How does redox imbalance in astrocytes lead to changes in neuronal resting voltage and neuronal activity? One possibility is that redox imbalance in astrocytes could impair astrocyte regulation of ion concentrations in the extracellular space, both through the activity of ion channels and transporters and by changing the volume of the extracellular space (*55*). Such changes in extracellular ion concentrations would directly alter neuronal resting membrane voltage. Astrocytes could also release trophic factors and signaling molecules that can influence the expression of neuronal ion channels and transporters (*56*). To determine how redox imbalance in astrocytes drives changes in neuronal activity, we performed RNA-Seq on cortical astrocytes isolated from four groups of animals: Con^Astro^, cKO^Astro^, SFN-fed Con^Astro^, and SFN-fed cKO^Astro^ (**Fig. 5A, S12A**). *Aldh7a1* and *Slc1a3* were significantly different between astrocytes isolated from cKO^Astro^ vs. Con^Astro^ mice as well as SFN-fed cKO^Astro^ vs. SFN-fed Con^Astro^ (**Fig. S12B; Table S1**), reasonably reflecting the animals’ genotype (*Aldh7a1*^*Flox/Flox*^; *Slc1a3*^CreER/+^). However, similar to the RNA-seq data from KO^Global^ brain tissue (**Fig. 2B**), only six other genes were significantly different between cKO^Astro^ and Con^Astro^ mice (*Acer2, Vsig2, Esr1, Foxd1, Echdc3*, and *A630020A06*). We therefore performed pathway analysis by GSEA to identify significant gene sets (pathways) potentially implicated in the SFN rescue of neuronal hypoactivity caused by astrocyte ALDH7A1 deficiency (**Fig. 5B; Table S1**). We focused on pathways with the following characteristics: 1) differentially expressed between untreated cKO^Astro^ and untreated Con^Astro^ astrocytes, 2) rescued in SFN-fed cKO^Astro^ astrocytes (i.e. no difference between SFN-fed cKO^Astro^ and untreated Con^Astro^ astrocytes), and 3) dissimilarly affected by SFN in Con^Astro^ vs. cKO^Astro^ astrocytes. Among the 262 pathways that fit these criteria (**Fig. 5C**), we identified 12 pathways related to ion channels and ion transporters as strong candidate mediators of reduced neuronal resting voltage (**Fig. 5D)**. Based on this bioinformatic analysis, we developed a working hypothesis that changes in astrocyte ion transporters and ion channels influence local extracellular ion concentrations, which in turn leads to prelimbic L5 neuronal hypoactivity in cKO^Astro^ mice.

**Fig. 5.**
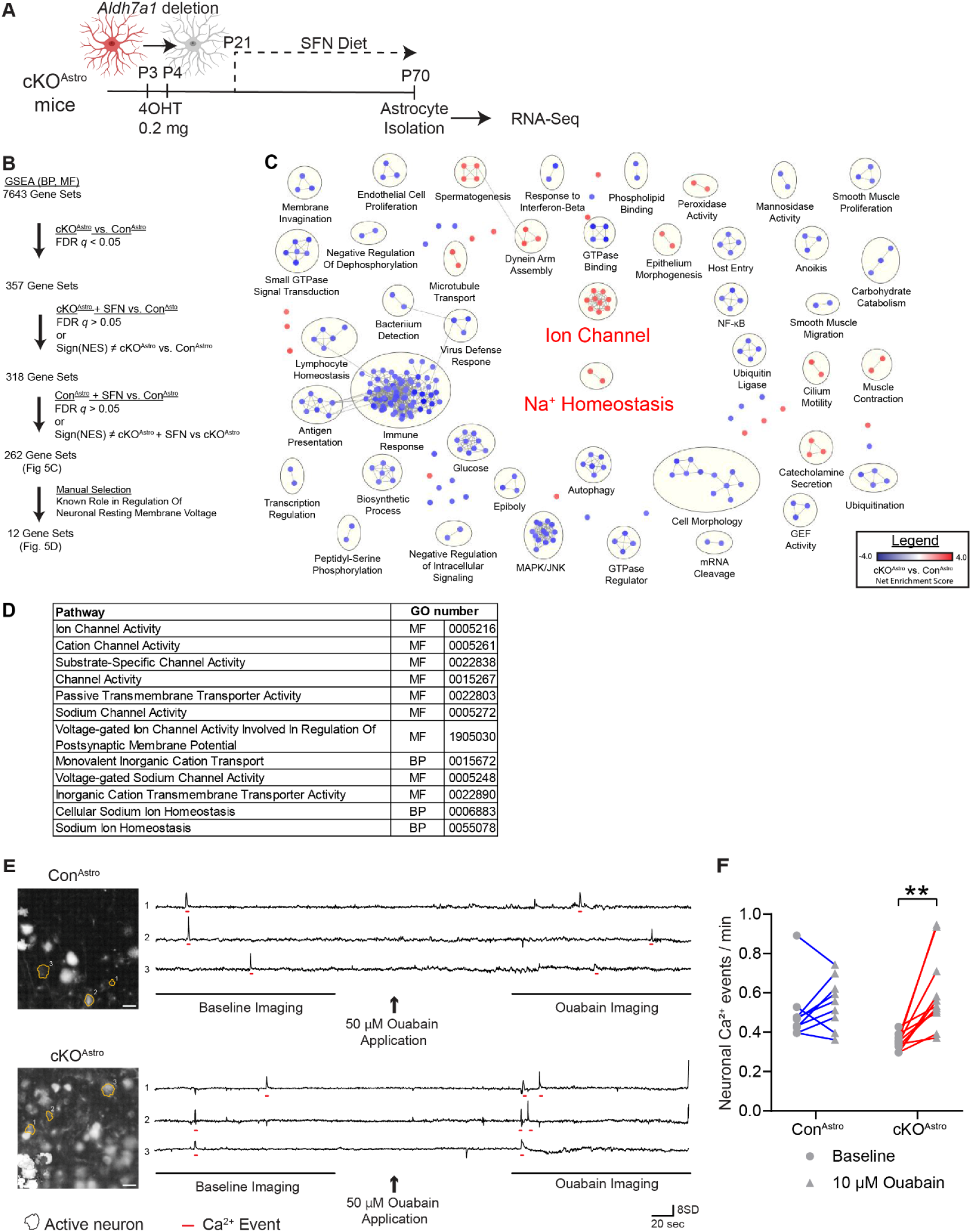
SFN ameliorates expression of ion channel and ion transporter gene pathways in cKO^Astro^ astrocytes. (A) Schematic of RNA-Sequencing of cortical astrocytes isolated from cKO^Astro^, SFN-treated cKO^Astro^ mice (cKO^Astro^ + SFN), control (Con^Astro^), and Con^Astro^ + SFN mice (B) Flow chart of post-hoc analysis of gene set enrichment analysis (GSEA) of astrocytes isolated from cKO^Astro^, cKO^Astro^ + SFN, Con^Astro^, and Con^Astro^ + SFN mice. From an initial list of 7643 affected gene sets, we narrowed down to 262 gene sets whose expression change patterns are suggestive of a genotype-specific rescue by SFN. (C) Visualization of the 262 gene sets identified in (B). Gene sets are represented as circles, color-coded based on their normalized enrichment score in cKO^Astro^ vs. Con^Astro^ astrocytes. Grey lines indicate gene sets with overlapping genes. Groups of gene sets with a high percentage of overlapping genes are indicated by yellow circles and labeled by biological theme. We manually highlighted two biological themes consisting of 12 gene sets potentially related to lower resting membrane voltage in neurons: ion channels and Na^+^ homeostasis. (D) Table of 12 gene sets potentially related to lower resting membrane voltage in neurons: ion channels and Na^+^ homeostasis. (E) Representative images (left) and traces (right) of GCaMP6f fluorescence in Con^Astro^ and cKO^Astro^ mice. Numbered orange outlines indicate neurons shown in traces. Events are indicated by red underlines. Time periods used for analysis are underlined in black. Arrows indicate application of 50 μM ouabain. Scale bars, 20 μm. (F) Quantification of prelimbic L5 pyramidal neuron Ca^2+^ event frequency in Con^Astro^ and cKO^Astro^ mice at baseline and in the presence of 50μM ouabain. Data represent individual brain slices, at baseline and after ouabain application (2-way repeated measures ANOVA with Holm-Sidak post-hoc tests: *n* = 9 brain sections. \*\**P* < 0.01).

It is known that astrocytes can affect neuronal activity by regulating extracellular potassium concentration [K^+^] through at least two mechanisms: active transport via Na^+^/K^+^ ATPases (*57*), or passive transport via two-pore K^+^ channels (*58*) and inward-rectifying K^+^ channels (specifically K_ir4.1_) (*55*). Within the “core enrichment” of the aforementioned 12 pathways (i.e. the subset of genes within those pathways that most contribute to the pathways’ significance; **Table S1**), we found genes encoding several subunits of Na^+^/K^+^ ATPases (*Atp1a1, Atp1a2, Atp1a3, Atp1b1, Atp1b2*, and *Atp1b3*) and genes encoding two-pore K^+^ channels (*Kcnk1, Kcnk2, Kcnk3, Kcnk4, Kcnk5, Kcnk7, Kcnk12*, and *Kcnk13*), but not K_ir4.1_ (*Kcnj10*). Since K_ir4.1_ gene expression was not significantly changed and two-pore K^+^ channels are challenging to specifically target by pharmacology, we focused on Na^+^/K^+^-ATPases for a further functional validation. To test whether astrocytic changes in Na^+^/K^+^ ATPases underlie the prelimbic L5 neuron hypoactivity, we applied a pharmacological inhibitor of Na^+^/K^+^ ATPases (50 μM ouabain) to acute brain slices and measured the effect on the prelimbic L5 pyramidal neuron Ca^2+^ event frequency (**Fig 5E-F**). Our hypothesis was that blocking Na^+^/K^+^ ATPases would rescue the neuronal hypoactivity. Consistent with prior experiments (**Fig. 4**), prelimbic L5 pyramidal neuron Ca^2+^ event frequency was significantly lower in cKO^Astro^ mice compared with that in Con^Astro^ mice. Within 3-6 min after the application of ouabain, Ca^2+^ event frequency in cKO^Astro^ neurons significantly increased, reaching the levels in Con^Astro^ mice, whereas Ca^2+^ event frequency in Con^Astro^ mice neurons did not significantly increase in this time frame, suggesting that excess astrocytic Na^+^/K^+^ ATPase activity, and associated reductions in extracellular [K^+^], are responsible for the prelimbic L5 pyramidal neuron hypoactivity. These data provide physiological validation of the bioinformatic inferences from cKO^Astro^ astrocyte gene expression data.

Lastly, we sought to determine whether the identified changes of ion transporter and ion channel genes in cKO^Astro^ astrocyte were under the influence of astrocyte redox status and SFN. To address this multifactorial question, we used the Causal Inference Engine (CIE) analysis platform (*59*), a bioinformatic approach to infer the activity of transcriptional regulators by comparing gene expression patterns of ion channel and ion transporter genes in cKO^Astro^ astrocytes to the ChIP-Seq Atlas, a publicly available database of transcription factor-gene regulatory networks (**Fig. S12C**). We restricted our analysis to the 432 ion channel and ion transporter genes that were identified by GSEA as the most significantly changed in the in cKO^Astro^ astrocytes (the “core enrichment” of the 12 ion channel and ion transporter pathways; **Table S1**). We identified transcription factors with significant differences in predicted activation in untreated cKO^Astro^ vs. untreated Con^Astro^ astrocytes, but not in SFN-fed cKO^Astro^ or SFN-fed Con^Astro^ astrocytes vs. untreated Con^Astro^ astrocytes (**Fig. S12D**). This list included ZBTB17, ZEB1, BRD2, SMAD1, FOXF2, GREB1, TET2, and GATA4, several of which are known to be redox-sensitive transcriptional regulators (*60-63*). Taken together, these bioinformatic analyses indicated that astrocyte ion channel and ion transporter gene changes in cKO^Astro^ mice are, at least in part, downstream of altered astrocyte redox homeostasis. Accordingly, an SFN-driven restoration of redox homeostasis was likely to provide the rescue effect.

## Discussion

The present study includes three major findings with implications for both basic and translational neuroscience. First, using animal models for a monogenic epilepsy, PDE, we dissociate two distinct brain pathophysiologies caused by liver-astrocyte pleiotropy: seizures are caused by hepatic ALDH7A1 depletion whereas affective behavior is caused by astrocytic ALDH7A1 depletion. Second, using cell type-specific manipulations, we define an astrocyte-initiated mechanism underlying affective behavior: ALDH7A1-dependent redox dysfunction in astrocytes causes prelimbic L5 pyramidal neuron hypoactivity linked to Na^+^/K^+^-ATPase-dependent reductions in extracellular [K^+^] that reduce neuronal resting membrane voltage. Lastly, we demonstrate that this mechanism in astrocytes is therapeutically actionable: NRF2 activation in astrocytes by viral and dietary approaches restores prelimbic activity and normalizes affective behavior, providing an opportunity for pathway-specific, cell type-targeted therapies orthogonal to antiseizure therapy.

Monogenic diseases can provide insight into the etiology of complex brain disorders, but many display pleiotropy across tissues and cell types, motivating cell-type specific mechanistic investigations. This strategy has been informative for neurodevelopmental disorders such as tuberous sclerosis (*TSC1/2* mutations), Rett’s syndrome (*MECP2* mutations), and Fragile-X syndrome (*FMR1* mutations) where both astrocytes and neurons shape distinct aspects of disease pathology (*64-66*). Our findings suggest that it is also important to consider gene function outside the CNS, including how peripheral pathologies may impact the brain. Glial cells share several functional specializations with peripheral cell types, raising the possibility that certain symptom clusters in monogenic conditions reflect contributions from both CNS and peripheral perturbations. For example, both astrocytes and hepatocytes have specialized roles in performing metabolic and detoxification functions for the brain and for the body, resulting in a large overlap between astrocyte-enriched and hepatocyte-enriched genes (e.g. *ALDH1L1, APOE, SOD1, GLUL, FABP7, CLU, GCLM, C3*) (*67, 68*). In the case of *ALDH7A1* mutations, this shared biology is reflected in a combination of hepatic abnormalities that reduce seizure threshold, reminiscent of established mechanisms of hepatic-induced seizures (*69-72*), and astrocyte-dependent changes in prelimbic neuronal activity and affective behavior. Notably, ALDH7A1 ablation in the liver and in astrocytes may influence different brain regions in different ways, which has parallels to patterns of focal hypo- and hyperactivity reported in epilepsy (*73-76*).

The mechanism linking astrocyte ALDH7A1 dysfunction with affective behavior aligns with emerging evidence that astrocytic regulation of neuronal activity plays a central role in mood disorders, particularly through K^+^ homeostasis. Extracellular [K^+^] impacts neuronal activity by setting the neuronal membrane potential and is controlled by astrocytes via Kir4.1 and Na^+^/K^+^-ATPases (*77, 78*). Both pathways have been implicated in depression (*79, 80*). Kir4.1 protein levels are upregulated in the postmortem parietal cortex from patients with major depressive disorder (MDD) (*81*) and astrocyte-specific manipulation of Kir4.1 in the lateral habenula alters ketamine-responsive affective behaviors in rodent models (*82*). Likewise, Na^+^/K^+^-ATPase subunit genes have decreased expression in MDD and bipolar disorder (BD) patients (*83, 84*) and are genetically linked to BD (*84*). ATP1A3 haploinsufficiency in mice also increases vulnerability to stress-induced depression-like phenotypes (*85*). These findings converge with our data showing that astrocyte Na^+^/K^+^-ATPases in the prelimbic cortex can modulate affective behavior by reducing extracellular [K^+^] and neuronal activity. However, a limitation of our pharmacological approach is that ouabain, while preferentially inhibiting astrocytic ATPases at the concentration used, may also weakly affect neuronal ATPases (*86*). More broadly, astrocyte morphology, gap-junction coupling, and Ca^2+^ signaling are all altered by stress and antidepressant treatment (*87-89*), suggesting that redox- and activity-dependent astrocyte-neuron interactions could represent a common pathophysiological pathway in mood disorders, distinct from seizure pathophysiology.

Future work is needed to more fully understand how ALDH7A1 deficiency in the liver elicits seizure pathology in the brain. Current models suggest that ALDH7A1 dysfunction leads to build up of metabolic intermediates that inactivate vitamin B6 (i.e. pyridoxine), which then inhibits over 100 enzymatic reactions in the brain that require vitamin B6 cofactor activity (*8*). The respective contributions of liver ALDH7A1 and astrocyte ALDH7A1 to vitamin B6 activity in the brain remain unknown. KO^Global^ mice do not exhibit visually detectable spontaneous seizures, which is ideal for distinguishing the direct effects of ALDH7A1 from secondary pathologies resulting from seizures, but phenotypically different from humans with PDE who suffer spontaneous seizures. Work from other murine models of ALDH7A1 ablation suggest that this apparent inter-species difference is largely due to diet rather than lack of homology – ALDH7A1 knockout mice don’t have spontaneous seizures on a standard lab mouse diet but exhibit spontaneous seizures when dietary lysine is increased or dietary pyridoxine is decreased (*30, 97*). Likewise, dietary changes in lysine and pyridoxine consumption can prevent spontaneous seizures in humans with PDE (*98*). In the current study, we use PTZ-induced seizures as a readout of seizure risk, which is reduced in KO^Global^ mice and responsive to pyridoxine supplementation, suggesting it is an effective readout of the brain metabolic state underlying seizures in humans that can be shifted up or down through dietary modifications of lysine and pyridoxine but relatively insensitive to astrocyte ALDH7A1 ablation.

Our work provides preclinical evidence that an indirect antioxidant approach such as dietary SFN may be an effective supplemental therapy for treating psychiatric symptoms in PDE patients in combination with anti-seizure therapies. Current clinical approaches, designed to combat metabolic deficiencies in vitamin B6 elicited by impairments in the lysine degradation pathway, rely on pyridoxine supplementation which is effective in controlling seizures but cannot treat mental/cognitive functions in PDE patients (*46*). Our data suggest that the selective effectiveness of pyridoxine is because pyridoxine treats the hepatic ALDH7A1 pathophysiology underlying seizures, but is ineffective in treating astrocyte ALDH7A1 pathophysiology underlying affective behavior. By contrast, dietary SFN, which targets astrocyte redox dyshomeostasis, is effective in normalizing affective behavior, but not seizures. Although there are challenges to modeling human affective behavior in rodents, the effectiveness of SFN in normalizing two prelimbic-associated behaviors with a similar dimension from a neuropsychology viewpoint suggests its potential utility to humans living with mood disorders. SFN is a promising lead compound since it is a brain-penetrant, naturally-occurring chemical safely tolerated in humans, already in clinical trials for psychiatric disorders including mood disorders and autism spectrum disorder (*90-94*). However, mechanism-driven trials of SFN are still pending (*95, 96*). In parallel to pursuing the translational potential of SFN itself, further mechanism-driven drug screening using SFN as a lead compound will also be important to establish novel therapeutic strategies. This study may therefore serve as a starting point for mechanism-driven treatment of psychiatric symptoms in PDE, separately from seizures, and a model for uncovering the pathophysiology of psychiatric symptoms in other epilepsy cases. To extend beyond PDE, future studies should incorporate astrocyte-associated biomarkers including GSH, prefrontal activity patterns, and assays of extracellular [K^+^]. Importantly, our data show that this pathway of astrocyte-elicited affective behavior is reversible, highlighting a broadly addressable astrocyte-driven pathway that can modulate affective behavior independent of seizures, a principle that extends beyond PDE.

## Materials and Methods

### Study design

Unless otherwise specified, experiments were performed in adult mice 10-16 weeks of age. All animals were healthy and were not immune compromised. Expression analyses were performed in C57BL/6NTac animals. Ibotenic acid and PALE experiments were performed in C57BL/6J mice. *Aldh7a1* ^*flox/flox*^ mice were generated and maintained on a C57BL/6NTac background (see Generation of *Aldh7a1* floxed mice section below). All additional transgenic lines were obtained on a C57BL/6 background and crossed with *Aldh7a1* ^*flox/flox*^ mice for at least two generations. Generation and genotyping of Cre driver lines (*CMV*^Cre^ (*99*), BAC-*Slc1a3*^CreER^ (*100), Slc1a3*^CreER^ (*101*), *Emx1*^Cre^ (*43*), *Alb1*^*Cre*^ (*102)*) and reporter lines (*R26*^lsl-mGCaMP3^ (*49*), *Slc1a2*^eGFP^ (*103*)) have been previously described. To generate KO^Global^ mice, *Aldh7a1* ^*flox/flox*^ mice were crossed with *CMV-Cre* mice and offspring were interbred to obtain *Aldh7a1*^−/–^; *CMV-Cre*^−^ mice (KO^Global^ mice). *Aldh7a1* ^*flox/flox*^ mice were crossed with *Slc1a3*^CreER/+^ mice to generate cKO^Astro^ mice. *Aldh7a1* ^*flox/flox*^ mice were crossed with *Emx1*^Cre^ mice to generate cKO^Emx1^ mice. *Aldh7a1*^*flox/flox*^ mice were crossed with *Alb1*^Cre^ mice to generate cKO^Liver^ mice. *Aldh7a1* ^*flox/flox*^ mice were crossed with BAC*-Slc1a3*^CreER^ mice to generate cKO^Astro(Mosaic)^ mice. Experimental cKO^Astro(Mosaic)^ and cKO^Astro^ mice were injected with 4-hydroxytamoxifen (see Tamoxifen section below) for induction of the Cre-loxP system from P3-P4 before sexing and genotyping. Genotyping was performed before weaning and reconfirmed post-mortem. For cKO^Astro^ and cKO^Liver^ mice, tamoxifen injected *Aldh7a1* ^*flox/flox*^ Cre-negative littermates were used as controls (Con^Astro^ and Con^Liver^). Both male and female mice were used for experiments to assess astrocyte function in KO^Global^, cKO^Astro(Mosaic)^, and cKO^Astro^ mice using immunofluorescence, biochemical assays, the mito-Grx1-roGFP2 ratiometric sensor, astrocyte calcium imaging, and neuronal calcium imaging in mice injected with astrocyte-targeting AAVs. Male mice were used for transcriptomic, electrophysiological, neuronal calcium imaging in mice treated with SFN, and behavioral experiments using the KO^Global^, cKO^Emx1^, cKO^Astro^, cKO^Liver^, and ibotenic acid models. Animals were group housed after weaning and maintained on a 12 h light/dark cycle with food and water provided *ad libitum*. For studies involving the pyridoxine diet or the SFN diet (see Pyridoxine diet and Sulforaphane diet sections below), cages were randomly assigned to receive standard diet or the experimental diet (pyridoxine or SFN). With the exception of behavioral studies, animals were never involved in previous procedures or studies. For behavioral experiments, animals were sequentially tested on open field test, forced swim test, and/or the seizure threshold test. Sucrose splash test behavior was assessed in separate cohorts of mice. All mice were used in accordance with Johns Hopkins University School of Medicine Institutional Animal Care and Use Committee (IACUC) guidelines.

### Generation of Aldh7a1 floxed mice

*Aldh7a1* floxed mice containing a *loxP*-flanked allele of *Aldh7a1* were produced by Taconic Biosciences. The targeting strategy was based on the NM_138600.4 transcript. The targeting vector contained *loxP* sites located in non-conserved regions flanking a ∼2 kb genomic region containing exons 4 and 5 of the *Aldh7a1* gene. Positive selection markers flanked by FRT (Neomycin resistance – NeoR) and F3 (Puromycin resistance – PuroR) sites were inserted into intron 3 and 5 respectively. The targeting vector was generated using BAC clones from the C57BL/6J RPCIB-731 BAC library and transfected into the C57BL/6J ES cell line. Homologous recombinant clones were isolated by double positive selection. The *Aldh7a1* floxed allele was obtained afterwards by Flp-mediated removal of the selection markers. Cre mediated recombination of the *Aldh7a1* floxed allele results in deletion of exons 4-5 and generates a frameshift from exon 3 to exons 6-11 resulting in a premature stop codon in exon 6.

### Genotyping of Aldh7a1 floxed mice

Routine genotyping of *Aldh7a1* ^*flox*^, *Aldh7a1*^*+*^, and *Aldh7a1*^−^ alleles was performed by polymerase chain reaction using the following primers at a 2:1:1 ratio: 5’-TCATAGCAGAGCACCTGATACC-3’, 5’-AAAGGCTTTGCACCACTGTG-3’, 5’-CCTATTGTGAGGGACTTTACCC-3’. These primers amplify a 177 bp DNA fragment for the wild-type allele, a 396 bp fragment for the floxed allele, and a 303 bp fragment for the knockout allele. Gels were visualized using Quantity-One software (Bio-Rad).

### Tamoxifen injections

Recombinant deletion of floxed alleles in CreER mouse lines was induced by 2 days of subcutaneous injection of 0.2 mg of 4-hydroxytamoxifen (4-OHT; Sigma H7904, St. Louis, MO, USA), once per day at P3 and P4. 4-OHT was dissolved in ethanol at 20 mg/mL by sonication for long term storage at -80 °C. On the day of injection, 4-OHT/ethanol solution was added to corn oil (1:5 ratio), vortexed for 4 min, and spun on a speed-vac for 30 min to evaporate the ethanol before injection.

### Immunofluorescence

According to established protocol (*104, 105*), mice were perfused with 1X TBS or PBS followed by 4% paraformaldehyde (PFA); brains were removed and postfixed in 4% PFA overnight.

For immunofluorescent analysis of ALDH7A1 expression, brain sections (30-40 μm) were collected using a vibratome or embedded in OCT compound with 7.5% sucrose after incubating in 30% sucrose and collected using a cryostat. Antigen retrieval was performed by incubation with L.A.B. solution (Polysciences, Inc.) when necessary: 10 min for 30 μm sections, 25 min for 100 μm sections, 1 h for 250 μm sections. Sections were permeabilized in TBS supplemented with 0.3% Triton X-100, blocked in 5% serum overnight at 4°C with primary antibody, washed in TBS, incubated with AlexaFluor-conjugated secondary antibody + DAPI for 2 h RT, washed and mounted on Superfrost Plus slides using ProLong Gold antifade reagent.

For immunolabeling of acute brain slices used for astrocyte microdomain Ca^2+^ imaging experiments, sections were incubated in primary antibody for ∼40 h at 4 °C followed by ∼16 h in secondary antibody at 4 °C and mounted with Vectashield mounting media with DAPI (Vector Labs H-1200).

For immunofluorescent analysis of NRF2, c-FOS, and MAP2, antigen retrieval was performed on 40 μm thick sections by preincubating in 1× Tris-EDTA (pH 9.0) (Abcam) at 80°C for 40 min [NRF2, c-FOS] or HistoVT One (Nacalai, 06380-05) at 65-70ºC for 30 min [MAP2]. Sections were then permeabilized in PBS supplemented with 0.5% Triton X-100 and blocked for 1 h in 10% serum. Subsequently, sections were incubated overnight at 4°C with Alexa Fluor-conjugated primary antibodies. After incubation, sections were washed in PBS containing DAPI and mounted on Superfrost Plus slides using ProLong Gold Antifade Reagent. Immunofluorescent analysis was performed in the same way for tissue from mice with stereotactic injection of AAVs for neuronal Ca^2+^ imaging.

*Antibodies used:* rabbit anti-ALDH7A1 EP1935Y (Abcam 53278; 1:250), chicken anti-GFP (Abcam 13970; 1:2,000), mouse anti-ALDH1L1 (Millipore MABN495; 1:1,000), mouse anti-CC1 (Calbiochem OP80; 1:50), guinea pig anti-NG2 (William Stallcup, Burnham Institute; 1:1,000), goat anti-IBA1 (Novus NB100-1028; 1:250), mouse anti-NeuN (Millipore MAB377; 1:500), mouse anti-GFAP (Millipore MAB360; 1:400), mouse anti-8-oxo-dG (Trevigen 4354-MC-050; 1:250), rabbit anti-NRF2-AlexaFluor488 (Abcam 194984; 1:100), anti-NRF2-AlexaFluor647 (Abcam 194985; 1:100), rabbit anti-ALDH1L1/2-AlexaFluor568 (Abcam ab313067; 1:100), rat anti-GFP (Nacalai 04404; 1:1000), rabbit anti-mCherry (Cell Signaling Technology 43590; 1:500), mouse anti-mCherry (Proteintech 68088-1-Ig; 1:400), rat anti-RFP (Proteintech 5f8; 1:1000), rabbit anti-NeuN (Cell Signaling Technology 12943; 1:250) and AlexaFluor 405-, 488-, 568-, 647-conjugated secondary antibodies.

*Imaging and analysis*: Immunofluorescent images were collected on Zeiss 800 and Zeiss 880 confocal microscopes using ZEN Blue/Black software.

For ALDH7A1 expression analysis, we collected z-stacks at 20x in three fields of view per mouse in the prelimbic cortex and, for each cell-type marker, counted the percentage cells that co-expressed ALDH7A1. Representative tiled images were acquired with 10x objective lens.

For anti-NRF2 analysis, we collected single z-plane confocal images at 40x in three fields of view per mouse from the prelimbic cortex. Quantification of anti-NRF2 fluorescence intensity in astrocytes and non-astrocyte regions was performed using FIJI software. Images were manually thresholded using the ALDH1L1 channel to identify astrocyte cell bodies and the mean fluorescence intensity was measured both within the ALDH1L1-positive region (astrocytes) and the ALDH1L1-negative region (non-astrocytes).

### Immunohistochemistry with 3,3’-Diaminobenzidine (DAB) development

After fixation, heart, lung, liver, kidney, and small intestine were processed, embedded in paraffin, and cut at 10 μm thickness. Deparaffinization and antigen retrieval were done in Dewax and HIER buffer M (Richard-Allen Scientific LLC: TA-999-DHBM) at 97 °C for 20 min on PT Module (Thermo Scientific: A80400012). Staining steps by DAB development were performed on Lab Vision Autostainer 360 (Thermo Scientific: A80500024). The tissues were incubated in 3% hydrogen peroxidase solution for 15 min at room temperature (RT). After washing with tris buffered saline (TBS), samples were blocked in TBS with 5% normal goat serum for 1 h at RT. ALDH7A1 antibody (abcam: ab53278) was incubated in blocking buffer for 1 h at RT. Samples were washed in PBS for 5 min x 3, then biotinylated anti-rabbit IgG (H + L) (vector: BA-1100) in TBS was applied and incubated for 1 h at RT. Samples were washed in TBS 3 times, and the ABC reagent prepared as instructed by the manufacturer was applied and incubated for 1 h at RT. Samples were washed in TBS 3 times, and DAB (Vector: SK-4100) solution was applied and incubated for 3 min at RT as instructed by manufacturer. The color developed samples were counterstained with hematoxylin (Thermo Scientific: 7211). After dehydration and serial incubation in xylene, samples were cover-slipped. Images were taken by an Olympus BX51TF with a DP70 color camera in the Microscope Facility of the Johns Hopkins School of Medicine or by Canon EOS 50D.

### Western Blot

Cortical tissue samples were obtained from adult mice following cervical dislocation and cortical dissection, snap frozen in liquid nitrogen, and stored at -80 °C. “Rest of brain” samples in which the cortices and cerebellum had been removed were also collected in some cases. Protein lysates were prepared by homogenization in lysis buffer (RIPA + protease inhibitors; Roche). For western blots, 20 µl of sample containing 20 µg of total protein was added to each lane of a Novex 8% Tris-Glycine gel (Life technologies) and run at 100 V at room temperature followed by transfer to a PVDF membrane at 4°C. Membranes were blocked for 1h in 5% milk in PBS + 1% Tween-20, incubated in primary antibody overnight, washed in PBS + tween, incubated in secondary antibody (anti-mouse or anti-rabbit HRP; GE Healthcare) for 1 h at RT, washed in PBS + tween, incubated with enhanced chemiluminescent substrate (ECL, Pierce), exposed, and quantified using an ImageQuant LAS 4000 mini (GE) digital image acquisition system. Raw images were quantified in ImageJ. Rectangular regions of interest (ROIs) were drawn around selected bands and mean gray value was reported by the measure tool. In each experiment, the same size ROI was used for all bands for every antibody. Background signal was measured by moving the ROI to an empty area of the blot and subtracted from the target signal prior to normalization to loading control signal. Blots were cropped, and in some cases, levels were adjusted for figure clarity. In these cases, image processing was always uniformly applied to all bands for each antibody. Experimenters were blind to the identity of the samples until analysis was complete. Antibodies used: rabbit anti-ALDH7A1 EP1934Y (Abcam 68192; 1:5,000), mouse anti-ALDH1L1 (Millipore MABN495; 1:1,000), mouse anti-GAPDH (Santa Cruz sc-32233, 1:10,000).

### Animal behavior

According to established protocol (*106, 107*), behavioral testing was performed in cohorts of 8-15 male mice starting at 10-14 weeks of age with tests in the following order: open field test, forced swim test, and seizure threshold test. Sucrose splash test behavior was assessed in separate cohorts of mice. Experiments were performed at room temperature. Experimenters were blind to the genotype of the animals during testing and video analysis of the behaviors. Stopwatch+ (Center for Behavioral Neuroscience, Atlanta, GA) was used for video analysis of forced swim test and seizure threshold test. Grooming events were counted using https://spacebarclicker.org/ (*108*).

*Open field test*: open field locomotor activity was measured for 120 min in a novel open field box (40 cm x 40 cm; San Diego Instruments, San Diego, CA). Horizontal and vertical locomotor activities were automatically recorded by an infrared activity monitor (San Diego Instruments). Single beam breaks were analyzed within 10 min bins and are reported as “counts.”

*Forced swim test*: forced swim response was measured by placing the mice in a 5000 mL glass beaker half-filled with water. We quantified the percentage of time the mice were actively swimming during the first two minutes.

*Sucrose splash test*: sucrose splash test was conducted according to established protocol by applying a 10% sucrose solution to the dorsal coat of each mouse within its home cage (*38*). The viscous nature of the sucrose solution ensures its adherence to the mouse fur, thereby eliciting grooming. Following the application of sucrose, two parameters were manually recorded in a blinded manner over a 5-minute observation period: the latency to initiate grooming and the frequency of grooming events. Self-grooming was specifically defined as licking of the fur or stroking and scratching of the face and body. These measures served as indices of self-care and motivational behavior.

*Seizure threshold test*: Each mouse was weighed, administered a series of 3 doses of PTZ at 30 min intervals (30 mg/kg, 30 mg/kg, and 60 mg/kg; i.p., Sigma P6500), and monitored until 30 min following the last dose. Seizure severity was scored on a modified Racine scale (*109, 110*) and the EC_50_ of the dose-response curve was estimated by a nonlinear least-squares regression model.

### Pyridoxine supplementation

Mouse chow containing 125 mg/kg pyridoxine (Cat# D16032903) or a standard diet (open standard diet; cat# D11112201) was obtained from Research Diets (New Brunswick, NJ). KO^Global^ mice were fed either pyridoxine-supplemented chow or standard chow starting 14 days before behavioral experiments. The food in the cage feeder was replaced every 7 days with fresh chow that had been stored at 4 °C or colder. Each time the food was changed, 21 g of diet was added to each cage for every mouse housed in it.

### RNA-Sequencing of cortical tissue

Cortical tissue was dissected from KO^Global^ mice and RNA was extracted using the RNeasy kit (Qiagen). Total RNA was assessed for quality via Agilent bioanalyzer and Thermo nanodrop, and RNA libraries were prepared using Illumina TruSeq Stranded RNA LT Kit with Ribo-Zero Gold rRNA depletion. Barcoded libraries were quality-controlled using the Kappa PCR kit and pooled in equimolar ratios for subsequent cluster generation and sequencing on an Illumina HiSeq 2000 instrument to yield >50,000,000 paired end 100x100 bp tags for each sample. *Preprocessing and aligning RNA-Seq reads to reference genome*: FastQC (*111*) was used to check the quality of reads. Reads were aligned to the mouse genome (mm10) using STAR (*112*) with the following optional parameters: --runThreadN 8 -- outSAMtype BAM SortedByCoordinate --quantMode GeneCounts. Count from ReadsPerGene table generated from STAR was used for further analysis

#### Differential gene expression analysis

Normalization and differential expression analysis were performed using the R package deseq2 (*113*). Genes with false discovery rate (FDR) smaller than 0.05 were identified as differentially expressed genes (DEGs).

#### Pathway analysis

We performed gene set enrichment analysis (GSEA) (*114*) on Gene Ontology (GO) biological processes and molecular function terms to identify over-represented pathways. All genes, including DEGs and non-DEGs, ranked by statistical significance obtained from differential expression analysis were used as the query list. GSEAPreranked module was used for the analysis. Pathways with less than 5 genes or more than 1000 genes were excluded (options - set_max 1000 -set_min 5). Pathways with FDR smaller than 0.05 were identified as significantly over-represented sets.

#### Network visualization

Gene sets of interest were loaded into Cytoscape for visualization. The Enrichment Map plug-in (*45*) was used to identify gene sets with overlapping genes. The AutoAnnotate plug-in was used to group gene sets with overlapping genes into biological themes. Biological themes were manually labeled to reflect the common function of the gene sets within that biological theme.

### Redox biochemical assays

Levels of NAD^+^/NADH, NADP^+^/NADPH, and reduced/oxidized glutathione (GSH/GSSG) were measured using commercial kits following the manufacturer’s instructions.

*NAD*^*+*^*/NADH*: NAD^+^/NADH was measured using a colorimetric assay kit (K337-100 BioVision, Inc. Melpitas, CA). Mouse brains were harvested, washed in PBS, and the cortex was isolated. Tissue samples (∼20 mg) were homogenized and sonicated in extraction buffer, centrifuged at 14,000 rpm for 5 minutes at 4°C, and filtered through 10 kDa molecular weight cutoff filters (Cat# 1997 BioVision). 200 μL of each sample was heated at 60°C for 30 minutes to degrade NAD^+^, then cooled on ice. 50 μL of degraded sample, 50 μL of undegraded sample, and 50 μL of NADH standards were loaded into a 96-well plate, mixed with NAD cycling reagent to convert the remaining NAD^+^ to NADH, incubated at room temperature for 5 minutes, mixed with NADH developer, and incubated at room temperature for 2 h. The amount of NADH was determined using endpoint measurement of absorbance at 450 nm and comparing to the standard curve. Measurements from the degraded samples reflected levels of NADH whereas measurements from undegraded samples reflected the total combined level of NAD^+^ and NADH. NAD^+^ levels were determined by subtracting NADH levels from the combined level of NAD^+^ and NADH, and then used to compute the NAD^+^/NADH ratio.

*NADP*^*+*^*/NADPH*: NADP^+^/NADPH levels were measured using a fluorometric assay kit (MET-5031 Cell Biolabs, Inc.). Mouse brains were perfused with cold PBS, and the prefrontal cortex was isolated by coronally cutting the brain at the optic chiasm, removing the olfactory bulbs, and extracting the portion of the cerebral cortex anterior to the optic chiasm. Tissue samples (100 mg) were homogenized and sonicated in extraction buffer, then centrifuged at 14,000 rpm for 5 minutes at 4°C. The supernatant was filtered using a 10 kDa spin filter to remove proteins, and the flow-through was collected for analysis and stored at -80°C. For NADPH measurement, samples were treated with NaOH, incubated at 80°C for 60 minutes, neutralized with assay buffer, vortexed, and centrifuged. NADP^+^ measurement followed a similar procedure using HCl. Samples and standards (50 µL) were loaded into a 96-well plate, mixed with NADP cycling reagent, and incubated for 1–2 h at room temperature. Fluorescence was measured at an excitation of 530–570 nm and an emission of 590–600 nm. NADP+/NADPH concentrations were determined using a standard curve.

*GSH/GSSG*: GSH/GSSG levels were measured using a colorimetric Glutathione Assay Kit (Cayman Chemical, MI; #703002). Mouse brains were perfused with cold PBS, and the prefrontal cortex was isolated by coronally cutting the brain at the optic chiasm, removing the olfactory bulbs, and extracting the portion of the cerebral cortex anterior to the optic chiasm. Tissue samples (100 mg) were homogenized and sonicated in cold MES buffer, then centrifuged at 10,000 g for 15 minutes at 4°C. The supernatant was removed and deproteinated. Metaphosphoric acid (MPA) reagent was prepared by dissolving 5 g of MPA in 50 mL of water. Equal volumes of MPA reagent were added to samples, vortexed, incubated at room temperature for 5 minutes, and centrifuged at >2,000 g for at least 2 minutes. The supernatant was carefully collected, avoiding disturbance of the precipitate. Homogenized samples were diluted to 1/10 and 1/100 using half-diluted MES buffer with MPA reagent. A 1 M solution of 2-vinylpyridine in ethanol was prepared. Ten microliters of this solution were added per milliliter of deproteinated sample, vortexed, incubated at room temperature for 60 minutes, and assayed. For the assay, 50 µL of standard and 50 µL of sample were added to each well of a 96-well plate. The plate was covered, and an assay cocktail was prepared by mixing MES buffer, reconstituted cofactor mixture, reconstituted enzyme mixture, and water. After removing the plate cover, 150 µL of freshly prepared assay cocktail was added to each well, and the plate was incubated in the dark on an orbital shaker. GSH concentration was determined using the endpoint method, measuring absorbance at 405–414 nm after 25 minutes.

### Cloning and generation of adeno-associated viruses

All cloning was performed using the Gibson Assembly method. The pZac2.1 GfaABC1D-mito Grx1-roGFP2 plasmid was generated by cloning the mito Grx1-roGFP2 coding DNA sequence from pLPCX mito Grx1-roGFP2 (Addgene Plasmid #64977) into the pZac2.1 gfaABC1D-tdTomato (Addgene Plasmid #44332) vector. The pZac2.1-GfaABC1D-mCherry plasmid was generated by replacing the tdTomato sequence from pZac2.1 GfaABC1D-tdTomato (Addgene Plasmid#44332) with the mCherry sequence from pAAV-CaMKIIa-mCherry (Addgene Plasmid#114469). The pZac2.1-GfaABC1D-ALDH7A1-mCherry plasmid was generated by inserting the mouse ALDH7A1 CDS (NCBI RefSeq: NM_138600.4) into the pZac2.1 GfaABC1D-mCherry plasmid. The pZac2.1-GfaABC1D-Nrf2-mCherry plasmid was generated by replacing the ALDH7A1 CDS in pZac2.1-GfaABC1D-ALDH7A1-mCherry with the *Nrf2* CDS from AAV-CMV-Nrf2 (Addgene Plasmid#67636). pGfaABC1D-mCherry-*Keap1*-shRNA and pGfaABC1D-mCherry-scramble-shRNA were generated by inserting a *Keap1* shRNA sequence (5’-CCTGCAACTCGGTGATCAATTC-3’) or a non-targeting scrambled shRNA sequence (5’-GGCTCCCGCTGAATTGGAATCC-3’) into a pAAV (AAV2 ITR–based) transfer backbone containing a pUC/ColE1 origin, Ampicillin resistance, and WPRE. The *Keap1* shRNA sequence was designed and embedded in an miR-30 backbone using VectorBuilder (https://en.vectorbuilder.com/) based on a validated siRNA sequence(*115*). The sequences for all newly generated plasmid sequences were deposited and are publicly available at GenBank. AAV5 viral particles were packaged at 1x10^13^ GC/mL by Applied Biological Materials Inc. (Richmond, BC, Canada).

### Assessment of astrocyte redox using the mito-Grx1-roGFP2 ratiometric sensor

AAV5-GfaABC1D-mito-Grx1-roGFP2 was bilaterally injected in the prelimbic cortex by stereotax (AP +1.94, ML ±0.3, DV -1.5) to express mito-Grx1-roGFP2 in astrocytes. Following established protocols (*116*), mice were perfused 4 weeks after AAV injection with phosphate-buffered saline (PBS) containing 50 mM N-Ethylmaleimide (NEM), followed by 4% paraformaldehyde (PFA) with 50 mM NEM. The extracted brains were then immersed in 4% PFA with 50 mM NEM overnight. Subsequently, brains were sectioned into 40 µm thick slices using a vibratome (Leica VT1200S). Sections were treated with 0.5% Triton X-100 in PBS containing 1 mM TO-PRO-3 (Thermo Scientific #T3605) for 20 minutes, followed by three washes in PBS. Finally, sections were mounted on Superfrost Plus slides using ProLong Gold antifade reagent. Fluorescence images were acquired frame by frame using a Zeiss LSM800 confocal microscope. Emission was collected from 500-700 nm, with sequential excitation provided by 405 nm and 488 nm laser lines. For each brain slice, a minimum of two images were captured using a 63x objective. Raw image data from the 405 nm and 488 nm laser lines were exported as 16-bit TIFF files for subsequent analysis with ImageJ software. To remove background fluorescence pixels, lower thresholds were applied to the 488 nm images. After defining selections based on these thresholds, the signal intensities for both the 488 nm and 405 nm images were measured.

### Astrocyte acute slice calcium imaging

Astrocyte calcium signals from acute brain slices containing prelimbic cortex were prepared, acquired, and analyzed using CasCaDe MATLAB code as previously described(*49*). Mice were deeply anesthetized with isoflurane and decapitated using a guillotine. Their brains were dissected out and mounted on a vibratome (Leica VT100S) equipped with a sapphire blade. Cortical slices 250 μm thick were cut in ice-cold N-methyl-D-glucamine (NMDG)-based cutting solution containing (in mM): 135 NMDG, 1 KCl, 1.2 KH_2_PO_4_, 1.5 MgCl_2_, 0.5 CaCl_2_, 10 Dextrose, 20 Choline Bicarbonate, (pH 7.4). Cortical slices were then transferred to artificial cerebral spinal fluid (ACSF) containing (in mM): 119 NaCl, 2.5 KCl, 2.5 CaCl_2_, 1.3 MgCl_2_, 1 NaH_2_PO_4_, 26.2 NaHCO_3_, and 11 Dextrose (292-298 mOsm/L) and were maintained at 37 °C for 40 min, and at room temperature thereafter. Both NMDG solution and ACSF were bubbled continuously with 95% O_2_/5% CO_2_. All slice imaging experiments were performed at room temperature. Fluorescence changes arising from mGCaMP3 were recorded from individual cortical astrocytes using a Zeiss LSM 710 microscope with a 20x (NA 1.0) water immersion objective (Zeiss), using the 488 nm laser line. GCaMP3^+^ astrocytes with no GCaMP3^+^ neighbors were selected for imaging in at least 3 cortical sections from 4 separate mice. Only one cell was imaged in each slice. Slices were continuously superfused with ACSF bubbled with 95% O_2_/5% CO_2_. Astrocytes were imaged at a laser power of 29 μW, with no pixel averaging, high PMT gain (∼800), and with a pinhole size of 2.69 airy units, corresponding to 3.6 μm of z-depth. For each experimental session, individual astrocytes (∼3500-4900 μm^2^) contained within prelimbic cortex were imaged at the pixel depth of 8 bit with a resolution of 512 × 512 pixels. Individual imaging sessions consisted of 600 frames at a frame scan rate of 2.1 Hz (0.484 s/frame). Each imaging session consisted of 5 min imaging to control for photoactivation prior to baseline measurements, 5 min baseline measurements, 15 min bath application of 200 nmol tert-butyl hydrogen peroxide (tBHP; Luperox 458139 Sigma) to induce ROS, 5 min imaging after tBHP application. To compensate for minor drift in the XY plane, image stacks were post hoc registered using TurboReg (plugin for ImageJ) for automatic alignment of images. Astrocytes located on the surface of the slice, those that had large blood vessels passing through them, or exhibited image registration artifacts were excluded from the dataset. Following imaging, the relative position of the cell in the section was recorded and a low magnification image was taken to allow the cell to be mapped during post-hoc immunohistochemistry by morphology, laser position, and the relative positions of nearby GCaMP3^+^ cells. We used the CasCaDe algorithm to identify and extract kinetic information about individual microdomain activity as previously described (*49*). Experimenters were blinded to the expression of ALDH7A1 in each astrocyte during imaging and analysis.

### Flow cytometry

For flow cytometry experiments, KO^Global^ mice were interbred with *Slc1a2*^EGFP/+^ (*103*) mice which express EGFP in all astrocytes. On each day of analysis, we analyzed one KO^Global^; *Slc1a2*^EGFP/+^ and one *Slc1a2*^EGFP/+^ littermate control. Cortices were dissected, separated from the hippocampus, and dissociated with a papain based neural tissue dissociation kit (Milenyi Biotec) to obtain a single cell suspension. After incubation, cell suspensions were spun down, supernatant was removed, and cells were resuspended in neurobasal media supplemented by 1% bovine serum albumin. For measurement of reactive oxygen species, 1µL of CellROX Deep Red reagent was added to samples containing 500,000 cells in 1mL and incubated at 37 °C for 30 min. Samples were then run on a MoFlo Legacy (Beckman Coulter) flow cytometer to analyze levels of CellROX fluorescence intensity in each cell population. Astrocytes were identified based on the expression of EGFP. Single color controls were used for compensation. Propidium iodide was added to the samples to exclude dead cells. The mean fluorescent intensity was calculated using FlowJo software. The fold change in mean fluorescent intensity between KO^Global^; *Slc1a2*^EGFP/+^ and one *Slc1a2*^EGFP/+^ astrocytes was calculated for each day of analysis.

### Mitochondrial stress test

Primary cortical astrocyte cultures were prepared from litters of *Aldh7a1*^−/–^ pups and *Aldh7a1*^*+/+*^ pups between P5-P9 of age. Litters contained both sexes and cells from each litter were pooled. Cortices were dissected, separated from the hippocampus, and dissociated with a papain based neural tissue dissociation kit (Miltenyi Biotec). Astrocytes were purified by magnetic activated cell sorting (MACS) technology using ACSA2^+^ microbeads (Miltenyi Biotec). Cells were grown in T-75 flasks precoated with poly-L-lysine (15μg/mL; Sciencell) in DMEM/F12 Dulbecco’s Modified Eagle Media supplemented with 15% fetal bovine serum and 1% penicillin/streptavidin and incubated at 37 °C in 5% CO_2_. After 14 days *in vitro* (DIV) cells were replated in a 24 well dish at 40,000 cells/well. Experiments were performed 2 days after replating.

Measurements of oxygen consumption rate (OCR) in primary astrocytes were performed on a Seahorse XF24 Extracellular Flux analyzer (Seahorse Bioscience). Two days before the assay, primary astrocytes were seeded into XF24 24-well microplates. The day before the assay, the cells were then placed in a 37°C incubator without CO_2_. In addition, an XF sensor cartridge was hydrated with Seahorse Bioscience Calibrant in an XF Utility plate and placed in a 37 °C CO_2_-free incubator overnight for 12 h. The day of the assay, CO_2_-free Seahorse Base Media was combined with 10 mM glucose, 1 mM sodium pyruvate, and 1x GlutaMAX (Thermofisher) and brought to pH 7.4. The cells were washed with this media and incubated in it for 1 h in the CO_2_-free incubator. During this incubation, the hydrated cartridge was loaded with oligomycin (1 µM), FCCP (2 µM), and antimycin A (0.5 µM) + rotenone (0.5 µM). The loaded cartridge was placed onto the cell well plate and inserted into the XF24 analyzer for repeated measurement of OCR and extracellular acidification rate (ECAR) at baseline and after sequential application of the mitochondrial blockers to assess mitochondrial basal respiration, ATP production, maximal respiration capacity, and spare capacity. Measurement cycles consisted of 3 min mix, 2 min wait, 3 min measure. Empty cells spaced throughout the plate are used as negative controls. These experiments were also repeated with application of 200 µM TBHP after baseline measurements to determine the effect of TBHP on cellular metabolism.

### Sulforaphane supplementation

Mouse chow containing 600 ppm D,L-Sulforaphane (cat# D15102501) or a standard diet (open standard diet; cat# D11112201) was obtained from Research Diets (New Brunswick, NJ). The D,L-Sulforaphane used for the diet was sourced from Toronto Research Chemicals, Toronto. Unless otherwise specified, treated mice and littermate controls were placed on a sulforaphane supplemented diet or a standard diet at the time of weaning, at 3 weeks of age. Cages of knockout mice and control littermates were randomly assigned to receive the SFN diet or the standard diet. The food in the cage feeder was replaced every 7 days with fresh chow that had been stored at 4 °C or colder. Each time the food was changed, 21 g of diet was added to each cage for every mouse housed in it.

### Measurement of sulforaphane in chow

Pellets stored at either 4 °C or ∼23 °C for 0, 6, 14, or 42 days were comminuted into fine powder using a small Krups coffee mill and extracted into 10 volumes of a chilled mixture of equal parts of water, acetonitrile, dimethylformamide, and dimethylsulfoxide. Sulforaphane was measured in these extracts by two separate methods, direct chromatography (HPLC) and cyclocondensation of total isothiocyanates (both free- and conjugated sulforaphane in the feed are captured by the latter method). 1,2-Benzenedithiol was purchased from Alfa Aesar (Cat. No. L11253) and stored at -20 °C. The sulforaphane standard was purchased from LKT Laboratories (Cat. No. S8044). Empower 3 software (Waters) was used for data acquisition.

#### Direct Chromatography

Extracts were clarified by centrifugation, diluted at least 10-fold in HPLC mobile phase (5% vol:vol tetrahydrofuran in water) then applied to a C18 reverse phase Genesis Lightning, 4.6 x 50 mm, 3 µm HPLC column (Grace Davison Discovery Sciences Cat. No. FM5963E), which was equilibrated with 5% tetrahydrofuran in water and eluted isocratically at 2 mL/min. Elution was monitored at 240 nm (the approximate λ_max_ of sulforaphane). The retention time was determined by comparison with an authentic sulforaphane standard purchased from LKT Laboratories.

#### Cyclocondensation

Total levels of isothiocyanate equivalents (isothiocyanates plus their metabolites) were measured using the HPLC–based cyclocondensation assay reaction with 1,2-benzenedithiol (*117, 118*). Isothiocyanates are metabolized to dithiocarbamates *in vivo*, and the cyclocondensation assay detects both isothiocyanates and dithiocarbamates. Briefly, each 2 mL reaction mixture in a 4 mL glass vial contained 100 mM borate buffer (pH 9.2), 16.3 mM 1,2-benzenedithiol, in 50% methanol/50% water. The reaction mixture was incubated for 2 h at 65 °C and, after cooling to room temperature, was centrifuged at low speed to sediment insoluble materials. Aliquots of the supernatant (50 and 100 µL replicates) were analyzed by HPLC, on an ODS C18 Hypersil, 100 x 4.6 mm, 5 µm column (Thermo Scientific #30105-104630). Mobile phase was 80% methanol/20% water, flow rate 0.5 mL/min, and absorbance was monitored at 365 nm as described previously (*117, 118*).

### Pharmacological lesioning

Pharmacological lesioning was performed by stereotactic injection of ibotenic acid (Sigma-Aldrich Cat# I2765), an NMDA glutamate receptor agonist (*119*). Bilateral injections of 1 μL ibotenic acid solution (2.5 μg/μL in saline) or vehicle (saline) were performed in either the prelimbic cortex (AP +1.94, ML ±0.3, DV -1.5) or the cerebellum (AP -6.4, ML ±2.7, DV -1.5), 4 weeks prior to behavioral testing.

### Neuronal acute slice calcium imaging and analysis

Calcium imaging experiments in cKO^Astro^ mice and KO^Global^ mice treated with sulforaphane and experiments using ouabain were performed using an epifluorescent microscope:

Four weeks prior to acute slice preparation and imaging, mice were stereotactically injected in the prelimbic cortex (AP +1.94, ML ±0.3, DV -1.5) with AAV9 pENN.AAV.CamKII.GCaMP6f.WPRE.SV40 (Addgene #100834). On the day of imaging, mice were anesthetized with ether, perfused using cold N-methyl-D-glucamine (NMDG) based cutting solution containing (in mM): 135 NMDG, 1 KCl, 1.2 KH_2_PO_4_, 1.5 MgCl_2_, 0.5 CaCl_2_, 10 dextrose, 20 choline bicarbonate, (pH 7.4) as previously described (*49*). Brains were dissected out and mounted on a vibratome (Leica VT100S). Cortical slices 250 μm thick were cut in ice-cold N-methyl-D-glucamine (NMDG) based cutting solution. Cortical slices were then transferred to artificial cerebral spinal fluid (ACSF) containing (in mM): 119 NaCl, 2.5 KCl, 2.5 CaCl_2_, 1.3 MgCl_2_, 1 NaH_2_PO_4_, 26.2 NaHCO_3_, and 11 dextrose (292–298 mOsm/L) and were maintained at 37 °C for 40 min, and at room temperature thereafter. Both the NMDG solution and ACSF were bubbled continuously with 95% O2/5% CO2. Experiments were carried out at room temperature.

Fluorescence dynamics arising from GCaMP6^+^ excitatory neurons in prelimbic cortex were recorded using a fixed-stage microscope (BX61-WI, Olympus) with optical lens (40x-0.80, LumPLanFL N), 300 W Xenon lamp using GFP filter, high speed galvo mirror with filters (DG4-1015, Shutter Instrument Company), and infrared-sensitive CCD camera (iXon3, ANDOR Technology). For each brain section, one field of view (FOV) in the prelimbic cortex was imaged for 600 frames in ∼200 s. Images were processed with digital imaging software (MetaFluor® for Olympus and Metamorph Advanced Molecular Device). All these systems are on an anti-vibration floating table (Technical Management Corp.) and connected to a PC (Windows 7, Microsoft). For imaging analysis, raw videos were pre-processed by applying ×4 spatial down-sampling to reduce file size and processing time, but no temporal down-sampling was applied (*120*). To reliably deal with the large fluctuating background, we applied the CNMFe algorithm (https://github.com/zhoupc/CNMF_E) that is a recent adaptation of the CNMF algorithm (*121*), enabling us to identify individual neurons, obtain their fluorescent traces, and deconvolve fluorescence signals into neural activity. To define the initial spatial components, candidate seed pixels were manually selected from peak-to-noise (PNR) graphs of the FOV (*122*). Calcium events with z-scores < 8 or those that did not have a >0.5 AUC were excluded from analyses because events of this magnitude did not reliably retain transient, calcium-event characteristics across animals. All neurons with at least one detected event were used to calculate the mean Ca^2+^ event rate.

For experiments using ouabain, each slice was imaged continuously for 600 s. 50 μM ouabain (MilliporeSigma, O3125) was added by bath application at 260 s. Ca^2+^ events from 0-200 s were analyzed and used determine the baseline Ca^2+^ event rate. Ca^2+^ events from 400-600 s were analyzed and used determine the Ca^2+^ event rate in the presence of ouabain. All neurons with at least one detected event both at baseline and in the presence of ouabain were used to calculate the mean Ca^2+^ event rate for each slice, both at baseline and in the presence of ouabain. To determine the effect of ouabain on Ca^2+^ event rate in cKO^Astro^ neurons vs. Con^Astro^, we performed a repeated-measures 2-way ANOVA.

Calcium imaging experiments in cKO^Astro^ mice with stereotactic injection of AAVs were performed using a spinning disk confocal microscope:

Four weeks prior to acute slice preparation and imaging, mice were stereotactically injected in the prelimbic cortex (AP +1.94, ML ±0.3, DV -1.5) with AAV9 pENN.AAV.CamKII.GCaMP6f.WPRE.SV40 (Addgene #100834) co-injected with either: AAV5-GfaABC1D-ALDH7A1 (ALDH7A1 Rescue), AAV5-GfaABC1D-mCherry (control AAV), AAV5-GfaABC1D-Nrf2 together with AAV5-GfaABC1D-mCherry-Keap1 shRNA (NRF2 Rescue), or AAV5 pZac2.1-GfaABC1D-mCherry together with AAV5 pGfaABC1D-mCherry-scramble RNA (control scramble AAV). On the day of imaging, mice were anesthetized and perfused as described above. For confocal imaging, cortical slices were 300 μm thick and transferred to ACSF. ACSF for ALDH7A1 Rescue experiments contained (in mM): 120 NaCl, 3.5 KCl, 2.5 CaCl_2_, 1.3 MgCl_2_, 1 NaH_2_PO_4_, 26.2 NaHCO_3_, and 11 dextrose (295–305 mOsm/L). ACSF for NRF2 Rescue experiments contained (in mM): 126.0 NaCl, 3.5 KCl, 1.0 CaCl_2_, 0.5 MgCl_2_, 1.25, 10.0 dextrose, and 10.0 HEPES (295–305 mOsm/L).Otherwise, slices and solutions were maintained as described above.

Confocal imaging of GCaMP6 fluorescence dynamics was performed on a Leica confocal microscope with a Yokogawa Spinning Disk unit (CSU10) for ALDH7A1 Rescue AAV or on a Nikon Eclipse TiE fully motorized inverted confocal microscope with a Yokogawa Spinning Disk unit (CSU10) for NRF2 Rescue AAVs. All these systems were on an anti-vibration floating table (Technical Management Corp.) and connected to a PC (Windows 11, Microsoft). During imaging, sections were kept in circulating ACSF containing (in mM): 126.0 NaCl, 3.5 KCl, 1.0 CaCl_2_, 0.5 MgCl_2_, 1.25 NaH_2_PO_4_, 26.0 NaHCO3, and 10.0 dextrose (295–305 mOsm/L). For each brain section, one field of view (FOV) in the prelimbic cortex was imaged for 900 frames in ∼300 s. Images were processed with SlideBook® (Intelligent Imaging Innovations, Inc.) and ImageJ software. Raw Ca^2+^ imaging movies were pre-processed in ImageJ with a custom macro (GenerateHitImage.ijm, available upon request) that applied bleach correction (simple ratio, background = 100), generated a difference image between maximum- and average-intensity projections, and performed rolling-ball background subtraction (radius = 20) to highlight active neurons for subsequent trace extraction and deconvolution of fluorescence signals into neural activity. Initial spatial components were defined by manually selecting candidate seed pixels from PNR graphs of the field of view (*122*). Calcium events with z-scores >2 were included in analyses, and all neurons with at least one detected event were used to calculate mean Ca^2+^ event rate.

### Analysis of c-FOS^+^ cells after chemically-induced seizures

Mice were administered one dose of PTZ (40 mg/kg i.p.) and monitored for seizure behavior. 60 min after administration of PTZ, mice that exhibited stage 7 seizures and survived were perfused with PBS followed by 4% PFA; brains were removed and post-fixed in PFA overnight. After buffer replacement with 30% sucrose, the brains were embedded and frozen in OCT compound with 7.5% sucrose. 40 μm coronal brain sections were cut using a cryostat. Sections were permeabilized in PBS supplemented with 0.5% Triton X-100 and blocked for 1 h in 10% serum. Subsequently, sections were incubated overnight at 4°C with primary antibody (rabbit anti-c-Fos; Cell Signaling Technology #2250; 1:500), washed in PBS, and incubated at RT with Alexa Fluor-conjugated secondary antibodies for 2 h. After incubation, sections were washed in PBS containing DAPI and mounted on Superfrost Plus slides using ProLong Gold Antifade Reagent. Immunofluorescent images were collected on a Zeiss 800 confocal microscope using ZEN Blue software. Representative tiled images were acquired at 10x. For quantification of c-FOS^+^ cells, we collected 3 single z-plane confocal images per region per animal in layer 5 of the prelimbic cortex at 20x and in the dentate gyrus at 63x. Quantification of the percentage of c-FOS+ cells was performed using FIJI software by dividing by the total number of DAPI^+^ cells. DAPI^+^ and c-FOS^+^ cells in the prelimbic cortex were quantified using Analyze Particles in FIJI and in the dentate gyrus by manual counting.

### Acute slice electrophysiology

Mice were anesthetized with Euthasol and perfused intracardially with 15ml ice-cold cutting ACSF (described below). Brains were quickly removed and placed in ice-cold cutting ACSF. Coronal sections (300 µm) containing PFC were prepared in ice-cold cutting ACSF using a vibrating blade microtome (Leica VT1200). Right after cutting, slices were recovered for 10 minutes at 32°C degrees and then transferred to holding ACSF at room temperature. Cutting and recovery were performed with ACSF containing the sodium substitute NMDG (Ting et al., 2014) (in mM): 92 NMDG, 20 HEPES (pH 7.35), 25 glucose, 30 sodium bicarbonate, 1.2 sodium phosphate, 2.5 potassium chloride, 5 sodium ascorbate, 3 sodium pyruvate, 2 thiourea, 10 magnesium, 14 sulfate, 0.5 calcium chloride. ACSF used for holding slices prior to recording was identical, but contained 92 mM NaCl instead of NMDG and 1 mM MgCl_2_ and 2 mM CaCl_2_. ACSF used to superfuse slices during recording contained (in mM): 125 NaCl, 2.5 KCl, 1.25 NaH2PO_4_, 1 MgCl_2_, 2.4 CaCl_2_, 26 NaHCO_3_, and 11 glucose. All ACSF solutions were saturated with 95% O2 and 5% CO2. For recording, a single slice was transferred to a heated chamber (32 °C) and superfused with normal ACSF (2.5 ml/min) using a peristaltic pump (WPI). Visualization of neurons in L5 of PFC was achieved with an upright microscope equipped for differential interference contrast (DIC) microscopy (BX51WI, Olympus). Whole-cell patch-clamp recordings were made using a MultiClamp 700B amplifier (1 kHz low-pass Bessel filter and 10 kHz digitization) with pClamp 10.3 software (Molecular Devices).

Voltage-clamp recordings of sEPSCs were made using glass pipets with resistance 2-4 MOhms, filled with internal solution containing (in mM): 117 cesium methanesulfonate, 20 HEPES, 0.4 EGTA, 2.8 NaCl, 5 tetraethylammonium chloride, 2.5 Mg-ATP, and 0.25 Na-GTP, pH 7.2–7.3 and 290 mOsm. sIPSCs were recorded with internal solution containing (in mM): 136 CsCl, 4 NaCl, 1.1 EGTA, 10 HEPES, 0.2 CaCl_2_, 4 Mg-ATP, 0.3 Na-GTP, pH 7.3 and 290 mOsm. Input resistance was monitored on-line during recordings; cells with access resistance changes greater than 20% were excluded from analysis. Recordings were made at -70mV holding potential and sampled at 1.8 kHz. sEPSCs were pharmacologically isolated by having gabazine (1 μM) and APV (50 μM) present throughout the experiment. sIPSCs were pharmacologically isolated with DNQX (10 μM) and APV (50 μM) in the bath. 200-300 events per cell were analyzed, using a threshold of 2X the baseline noise. Analysis of sEPSCs and sIPSCs was performed off-line using the MiniAnalysis program (v 6.0, Synaptosoft) by experimenters blind to the genotype of the animals.

Current-clamp recordings were made in the absence of drugs with internal solution containing (in mM): 125 potassium gluconate, 0.05 EGTA, 10 HEPES, 6.27 KCl, 5 Na2-phosphocreatine, 4 Mg-ATP, and 0.3 Na-GTP, pH 7.3 and 285 mOsm. Rheobase was determined by injecting 2 ms steps of positive current (increasing in 25pA increments per sweep); values reflect size in pA of the first current injection that elicited an action potential. Number of spikes per current clamp step were calculated from action potentials elicited by a 1 s current injection (increasing in 25 pA increments per sweep).

### Postnatal astrocyte labeling by electroporation (PALE)

PALE is a medication of *in utero* electroporation to introduce plasmid constructs into astrocytes by shifting the age of electroporation to P0-P1 (*123*). To perform PALE, newborn P0 pups were anesthetized by hypothermia on ice for 4-5 min. A plasmid mixture containing pZac2.1-GfaABC1D-Nrf2-mcherry (1 µg/µL) and pZac2.1-GfaABC1D-mCherry-*Keap1* shRNA (1 µg/µL) was injected into the lateral ventricles using a glass micropipette made from a capillary tube (Narishige, Cat. #GD-1), followed by electroporation into the ventricular zone of the cortical plate. For electroporation, electrical pulses (40 V, 50 ms duration) were delivered four times at 950 ms intervals using an electroporator (Nepagene, Cat. #CUY21EDIT).

### RNA-Sequencing analysis of isolated astrocytes

Cortical astrocytes were acutely isolated from adult cortices by MACS cell separation. Briefly, cortical tissue samples were obtained from adult mice following saline perfusion and cortical dissection, homogenized using a papain dissociation kit (Miltenyi Biotec #130-107-677), incubated with Fc blocking antibody and magnetic anti-ACSA-2 microbeads (Miltenyi Biotec #130-097-679), bound to MACS LS columns (Miltenyi Biotec #130-042-201) placed on a magnetic stand, washed 3 times with 0.5% BSA buffer to remove unbound cells, and eluted for downstream application. RNA was extracted from isolated astrocytes by using the picopure RNA isolation kit (Applied Biosystems). Samples were checked for quality/quantity on Fragment Analyzer and Qubit prior to library prep (RQN > 7; 100-400 ng per sample). NEBNext Poly(A) Magnetic Isolation Module (NEB #E7490) and NEBNext Ultra II RNA Library Prep Kit for Illumina (Cat# E7775) were used to generate libraries. Paired-End Sequencing was performed using Illumina’s NovaSeq6000 S1 200 cycle kit. Preprocessing and aligning RNA-Seq reads to reference genome, differential gene expression analysis, pathway analysis, and network visualization Enrichment Map were performed as described above (see “RNA-Sequencing of cortical tissue”).

#### Upstream transcription factor analysis

Through GSEA analysis, we identified 12 GO terms related to ion channel and ion transporter activity (GO:0005216, GO:0005261; GO:0022838; GO:0015267; GO:002803; GO:0005272; GO:1905030; GO:0015672; GO:0005248; GO:0022890; GO:0006993; GO:0055078) as biologically relevant pathways whose disturbance caused by ALDH7A1 deficiency were rescued by SFN. We next conducted Causal Inference Engine (CIE) analysis (*59*) to identify up-stream regulators of core genes (genes that contribute to the leading-edge subset within the gene set) of the GO terms related to ion channel activity identified by GSEA. Specifically, ternary scoring statistic proposed by Chindelevitch *et al. (124*) was conducted to query ChIP-seq derived TF–gene interactions from ChIP-Atlas (*125*) and identify transcriptional regulators from differentially expressed genes (relaxed cutoff of p-value < 0.05 was used here) identified by DESeq2.

### Statistics

Representative immunofluorescent images shown were qualitatively similar across at least 3 animals. Individual data points are displayed for all experiments containing multiple data points. Data are reported as mean ± S.E.M. The exact value of *n* for each experiment is listed in the figure legends. For Western blot, ALDH7A1 cell-type expression, animal weight, behavior, RNA-Seq, and flow cytometry, *n* represents the number of animals analyzed per condition. For measurements of whole cell recordings, experiments in cKO^Astro(Mosaic)^ mice, and spontaneous neuronal Ca^2+^ imaging, *n* represents the number of cells analyzed per condition, taken from at least 3 mice per condition. For mitochondrial stress test measurements, *n* represents the number of primary cultures analyzed, each of which contained 10 technical replicates. For analysis of neuronal Ca^2+^ imaging in response to ouabain, *n* represents the number of acute brain slices analyzed per condition. A specific randomization strategy was not used and no statistical computations were performed to determine sample sizes for most experiments. We used sample sizes consistent with other studies in the field. Statistical analyses were performed using GraphPad Prism 10 software. Comparisons of PTZ dose-response curves were calculated by the extra-sum-of-squares F test of EC_50_ values. *P*-values for two groups were calculated by two-tailed Student’s *t*-test and between three or more groups by one-way ANOVA followed by Tukey’s multiple comparison test or Dunnett’s post-hoc test. Comparisons between genotypes across two conditions were calculated by two-way ANOVA or mixed-model analysis followed by Holm-Sidak’s multiple comparison tests between genotypes. When multiple measurements were performed in the same animal or cell, we performed a repeated-measure two-way ANOVA followed by Holm-Sidak’s multiple comparison tests between genotypes. Non-parametric comparisons between 3 or more groups were made using Kruskal-Wallis test with Dunn’s post hoc test. 2-way non-parametric comparisons between genotypes across two conditions were made using multiple Mann-Whitney tests with Holm-Sidak correction for multiple comparisons. The statistical tests used to measure significance are indicated in each figure along with the corresponding significance level.

## Acknowledgments

We thank Ms. Maria Papapavlou, Mr. Saarang Deshpande, Mr. Hao Zhang, Ms. Kristina Wade, Ms. Katherine Stephenson, Ms. Thanh Hai Tran, Ms. Yunqing Wang, and Dr. Tomohide R. Sato for technical assistance, as well as Dr. Mikhail Pletnikov, Dr. Jeffrey Rothstein, and Dr. Heather Faust for helpful discussions. We also thank Ms. Yukiko Lema for assistance preparing the figures and Dr. Hoku West-Foyle for help with ImageJ macros used for Ca^2+^ imaging analysis. We are grateful to Dr. Jeffrey Rothstein for providing *Slc1a2*^EGFP^ mice, Dr. Magdalena Götz for providing *Slc1a3*^CreER^ mice, and Dr. Cagla Eroglu for sharing protocols and technical advice.

## Funding

National Institutes of Health grant MH-107730 (ASawa)

National Institutes of Health grant MH-105660 (ASawa)

National Institutes of Health grant MH-094268 (ASawa)

National Institutes of Health grant MH-136297 (ASawa)

Stanley (ASawa)

S-R/RUSK (ASawa)

NARSAD (ASawa)

Subsidies for Current Expenditures to Private Institutions of Higher Education from the Promotion and Mutual Aid Corporation for Private Schools of Japan (KI)

National Institutes of Health grant NS050274 (Multiphoton Imaging Core)

NIDA Intramural Research Program of the NIH (WX)

NSF grant no. 1232825 (WX)

NARSAD young investigator grant 24161 (AA)

Chica and Heinz Schaller Research Foundation grant (AA)

## Author contributions

Conceptualization and Methodology: TEF and ASawa

Formal analysis: TEF, ASaito, KY, WX, SI, HN, AA, BL, and KI

Investigation: TEF, ASaito, KY, WX, SI, HN, AA, AR, BL, LH, RS, SA, SS, TP, TC-P, DW, EC, JWF, H J-P, and KI

Supervision: DEB

Writing: TEF and ASawa

## Data and materials availability

All published reagents will be shared on an unrestricted basis. The data that support the findings of this study are available from the corresponding author upon reasonable request. The RNA-Seq expression data are available at NCBI Gene Expression Omnibus (GEO) under accession number GSEXXXX (GSE id will be provided when the paper is provisional accepted). The CasCaDe MATLAB code for analysis of microdomain Ca^2+^ events was previously described and is freely available (*49*). Further information and requests for resources and reagents should be directed to and will be fulfilled by the Lead Contact, Akira Sawa.

