## Supplementary figures for "Astrocyte redox imbalance underlies prelimbic neuronal hypoactivity and affective behavior in epilepsy"

**The PDF file includes:**

Fig. S1. Molecular validation and baseline behavior of ALDH7A1 knockout mice.

Fig. S2. ALDH7A1 expression is not heterogeneous across brain regions or sexes.

Fig. S3. ALDH7A1 deletion does not induce astrocyte reactivity or robust oxidative stress.

Fig. S4. Validation of sulforaphane diet.

Fig. S5. Behavioral analysis of sulforaphane diet.

Fig. S6. Pharmacological lesioning of prelimbic cortex impairs sucrose splash behavior.

Fig. S7. Reduced layer 5 pyramidal neuron activity in the prelimbic cortex of $KO^{Global}$ mice.

Fig. S8. $KO^{Global}$ mice have a reduced percentage of c-$FOS^{+}$ cells in the prelimbic cortex during seizures.

Fig. S9. No changes in spontaneous postsynaptic currents on layer 5 pyramidal neurons in the prelimbic cortex of $cKO^{Astro}$ or $KO^{Global}$ mice.

Fig. S10. Reduced resting membrane voltage and excitability of layer 5 pyramidal neurons in the prelimbic cortex of $KO^{Global}$ mice.

Fig. S11. Validation of ALDH7A1 and NRF2 Rescue constructs.

Fig. S12. Putative upstream regulators of ion channel and ion transporter gene changes in $cKO^{Astro}$ astrocytes include redox-sensitive transcription factors.

**The supplementary materials also include the following files:**

Table S1. Bioinformatic analysis of RNA sequencing results. (Excel file).

**Fig. S1**

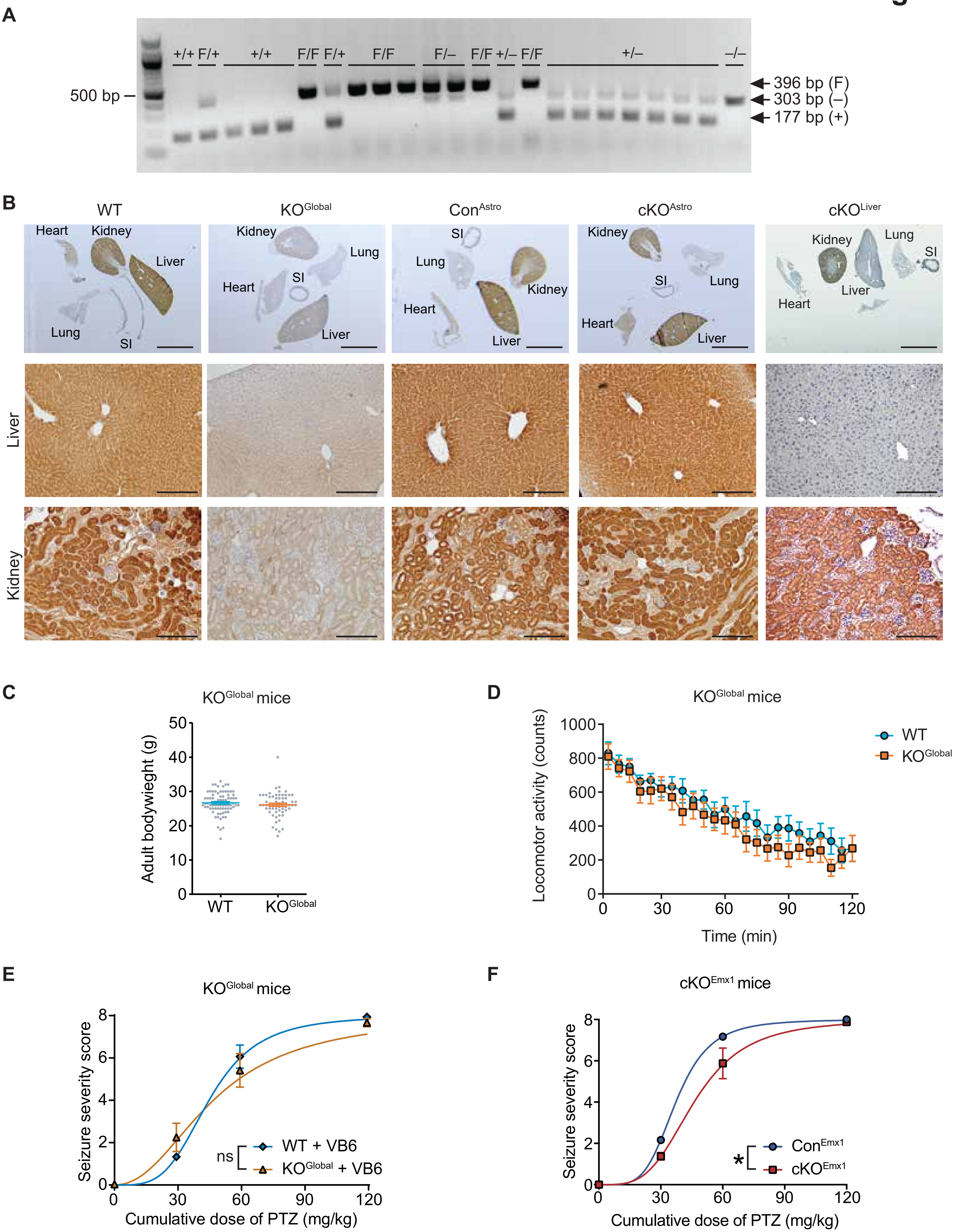

A
+/+ F/+ +/+ F/F F/+ F/F F/– F/F +/– F/F +/– –/–
500 bp
396 bp (F)
303 bp (–)
177 bp (+)
B
WT
KO^Global
Con^Astro
cKO^Astro
cKO^Liver
Heart
Kidney
Liver
Lung
SI
Liver
Kidney
C
KO^Global mice
Adult bodywieght (g)
WT
KO^Global
D
KO^Global mice
Locomotor activity (counts)
Time (min)
WT
KO^Global
E
KO^Global mice
Seizure severity score
Cumulative dose of PTZ (mg/kg)
ns
WT + VB6
KO^Global + VB6
F
cKO^Emx1 mice
Seizure severity score
Cumulative dose of PTZ (mg/kg)
*
Con^Emx1
cKO^Emx1

**Fig. S1. Molecular validation and baseline behavior of ALDH7A1 knockout mice.**

(A) Gel electrophoresis of polymerase chain reaction products from *Aldh7a1* floxed mice. Bands amplified from floxed (F), wild type (+), and knockout (–) alleles of *Aldh7a1* are indicated.
(B) Representative images of ALDH7A1 DAB staining in peripheral tissues (heart, kidney, liver, lung, and small intestine (SI)) in wild-type (WT), $KO^{Global}$, $Con^{Astro}$, $cKO^{Astro}$, and $cKO^{Liver}$ mice. Lower panels show high magnification images of liver and kidney. Upper panel scale bars, 5 mm. Lower panel scale bars, 200 μm.
(C) Quantification of adult bodyweight of $KO^{Global}$ and WT mice (Student's t-test: $n$ = 62-78 mice; $P > 0.05$)
(D) Quantification of locomotor activity of $KO^{Global}$ and WT mice during an open field test (2-way repeated measures ANOVA: $n$ = 11-14 mice; genotype effect: $P > 0.05$)
(E) Quantification of seizure severity in response to repeated doses of pentylenetetrazol (PTZ) during seizure threshold test in $KO^{Global}$ and WT mice on a diet supplemented with pyridoxine (VB6). Lines represent fitted dose-response curves (Extra sum-of-squares F test on dose-response $EC_{50}$: $n$ = 12-15 mice; ns: $P > 0.05$).
(F) Quantification of seizure severity in response to repeated doses of pentylenetetrazol (PTZ) during seizure threshold test in $cKO^{Emx1}$ and $Con^{Emx1}$ mice. Lines represent fitted dose-response curves (Extra sum-of-squares F test on dose-response $EC_{50}$: $n$ = 6-8 mice; ns: *$P < 0.05$).
All data represent mean ± S.E.M.

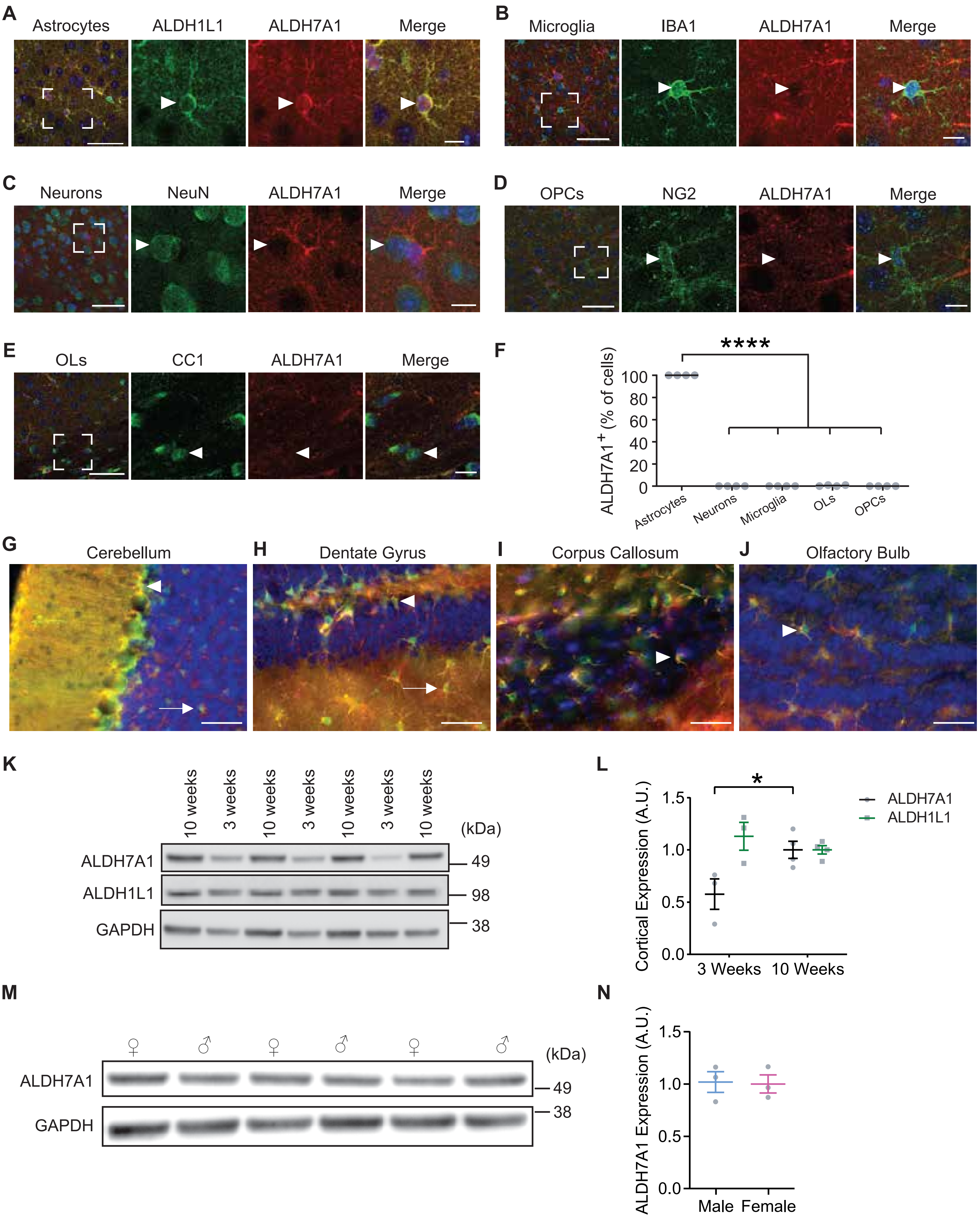
A
Astrocytes
ALDH1L1
ALDH7A1
Merge
B
Microglia
IBA1
ALDH7A1
Merge
C
Neurons
NeuN
ALDH7A1
Merge
D
OPCs
NG2
ALDH7A1
Merge
E
OLs
CC1
ALDH7A1
Merge
F
****
ALDH7A1+ (% of cells)
100
80
60
40
20
0
Astrocytes
Neurons
Microglia
OLs
OPCs
G
Cerebellum
H
Dentate Gyrus
I
Corpus Callosum
J
Olfactory Bulb
K
10 weeks
3 weeks
10 weeks
3 weeks
10 weeks
3 weeks
10 weeks
(kDa)
ALDH7A1
49
ALDH1L1
98
GAPDH
38
L
*
Cortical Expression (A.U.)
1.5
1.0
0.5
0.0
3 Weeks
10 Weeks
ALDH7A1
ALDH1L1
M
♀
♂
♀
♂
♀
♂
(kDa)
ALDH7A1
49
GAPDH
38
N
ALDH7A1 Expression (A.U.)
1.5
1.0
0.5
0.0
Male
Female

**Fig. S2. ALDH7A1 expression is not heterogeneous across brain regions or sexes.**

(A-E) Representative images of ALDH7A1 immunofluorescence in the cortex of adult wild-type (WT) mice, co-labeled with (A) astrocyte marker ALDH1L1, (B) microglia marker IBA1, (C) neuronal marker NeuN, (D) oligodendrocyte precursor cell (OPC) marker $NG2^{+}$, and (E) mature oligodendrocyte (OL) marker CC1 in adult mouse cortex. Boxed insets displayed at higher magnification in right three panels. Arrowheads indicate representative cells labeled by cell type markers. Scale bars: 50 µm. Inset scale bars: 10 µm.

(F) Quantification of the percentage of astrocytes (Aldh1l1), neurons (NeuN), microglia (Iba1), oligodendrocytes (CC1), and oligodendrocyte precursor cells (NG2) expressing ALDH7A1 in adult mouse cortex (1-way repeated measure ANOVA with Tukey post-hoc test: $n = 4$ mice; ****$P < 0.0001$).

(G-J) Representative images of ALDH7A1 immunofluorescence (red) in the (G) cerebellum, (H) dentate gyrus, (I) corpus callosum, and (J) olfactory bulb of adult *Slc1a2*$^{eGFP/+}$ reporter mice. Arrowheads indicate colocalization of ALDH7A1 with GFP (green) in Bergmann glia in (G), adult neural stem cells in (H), fibrous astrocytes in (I), and olfactory bulb astrocytes in (J). Arrows indicate colocalization of ALDH7A1 in velate astrocytes in (G) and hippocampal astrocytes in (H). Scale bars: 100 µm.

(K-L) ALDH7A1 proteins levels in homogenized cortical tissue at 3 weeks (3w) and 10 weeks (10w) of age. (K) Western blot images. GAPDH included as loading control and ALDH1L1 as an astrocyte cell-type marker (L) Quantification of ALDH7A1 and ALDH1L1 levels, normalized to GAPDH, at 3w vs. 10w (2-way repeated measures ANOVA with Holm-Sidak post-hoc test: $n = 3\text{-}4$ mice; *$P < 0.05$).

(M-N) Western blot of ALDH7A1 in adult cortical tissue of male (♂) and female (♀) mice. (M) Western blot image. (N) Quantification of ALDH7A1 levels normalized to GAPDH (Student's t-test: $n = 3$ mice; $P > 0.05$).

All data represent mean ± S.E.M.

### Fig. S3

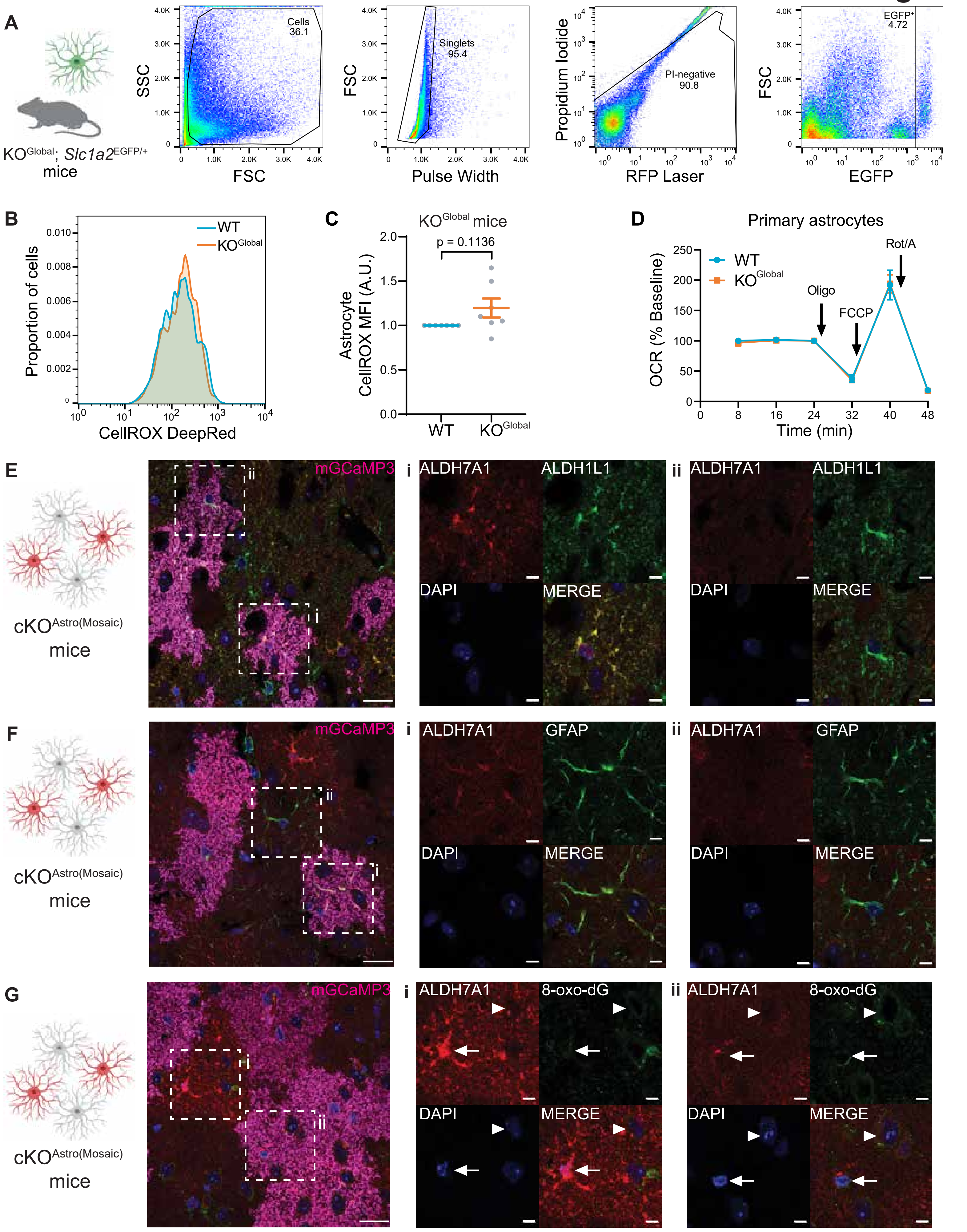
A
KO^Global; Slc1a2^EGFP/+ mice
SSC
FSC
Cells 36.1
Pulse Width
Singlets 95.4
Propidium Iodide
RFP Laser
PI-negative 90.8
EGFP
EGFP+ 4.72
B
Proportion of cells
CellROX DeepRed
WT
KO^Global
C
KO^Global mice
p = 0.1136
Astrocyte CellROX MFI (A.U.)
WT
KO^Global
D
Primary astrocytes
OCR (% Baseline)
Time (min)
Oligo
FCCP
Rot/A
E
cKO^Astro(Mosaic) mice
mGCaMP3
ALDH7A1
ALDH1L1
DAPI
MERGE
F
GFAP
G
8-oxo-dG

**Fig. S3. ALDH7A1 deletion does not induce astrocyte reactivity or robust oxidative stress.**

(A) Gating strategy for acutely isolated astrocytes from adult KO$^{Global}$; *Slc1a2*$^{eGFP/+}$ mice. After gating for size and singlets, propidium iodide (PI) was used to identify live cells. *Slc1a2*-EGFP was then used to identify astrocytes.

(B) Unit area histograms of CellROX DeepRed, a fluorescent indicator of reactive oxygen species, in live astrocytes from a representative pair of WT and KO$^{Global}$ littermates.

(C) Mean fluorescence intensity (MFI) of CellROX in live astrocytes in WT and KO$^{Global}$ mice. Data from each pair of WT and KO$^{Global}$ mice are normalized to WT levels. *(*1-sample t-test: $n$ = 7 mice; $P$ = 0.1136).

(D) Oxygen consumption rate (OCR) of primary astrocytes cultured from KO$^{Global}$ and WT mice during a Seahorse mitochondrial stress test. Arrows indicate when 1 μM oligomycin (an ATP synthase inhibitor), 2 μM FCCP (an uncoupling agent), and 0.5 μM Rotenone + 0.5 μM Antimycin (Rot/A; complex I and complex III inhibitors) were added (2-way repeated measures ANOVA with Holm-Sidak post-hoc tests between genotypes: $n$ = 3 separate culture experiments, $P > 0.05$).

(E-G) (Left) Schematic of ALDH7A1 depletion in cKO$^{Astro(Mosaic)}$ mice and (right) representative immunofluorescent images of ALDH7A1 (red) and mGCaMP3 (pseudocolored magenta) in cortex of adult cKO$^{Astro(Mosaic)}$ mice with (E) ALDH1L1, (F) GFAP, and (G) 8-oxo-dG. Scale bars: 20 μM. Insets depict high magnification images of an ALDH7A1$^{+}$ cell (i) and an ALDH7A1$^{-}$ cell (ii). In (G), nuclear 8-oxo-dG immunofluorescence is highlighted in the astrocyte (arrow) and a cell surrounded by astrocytic processes (arrowhead). Inset scale bars: 5 μM.

All data represent mean ± S.E.M.

#### Fig. S4

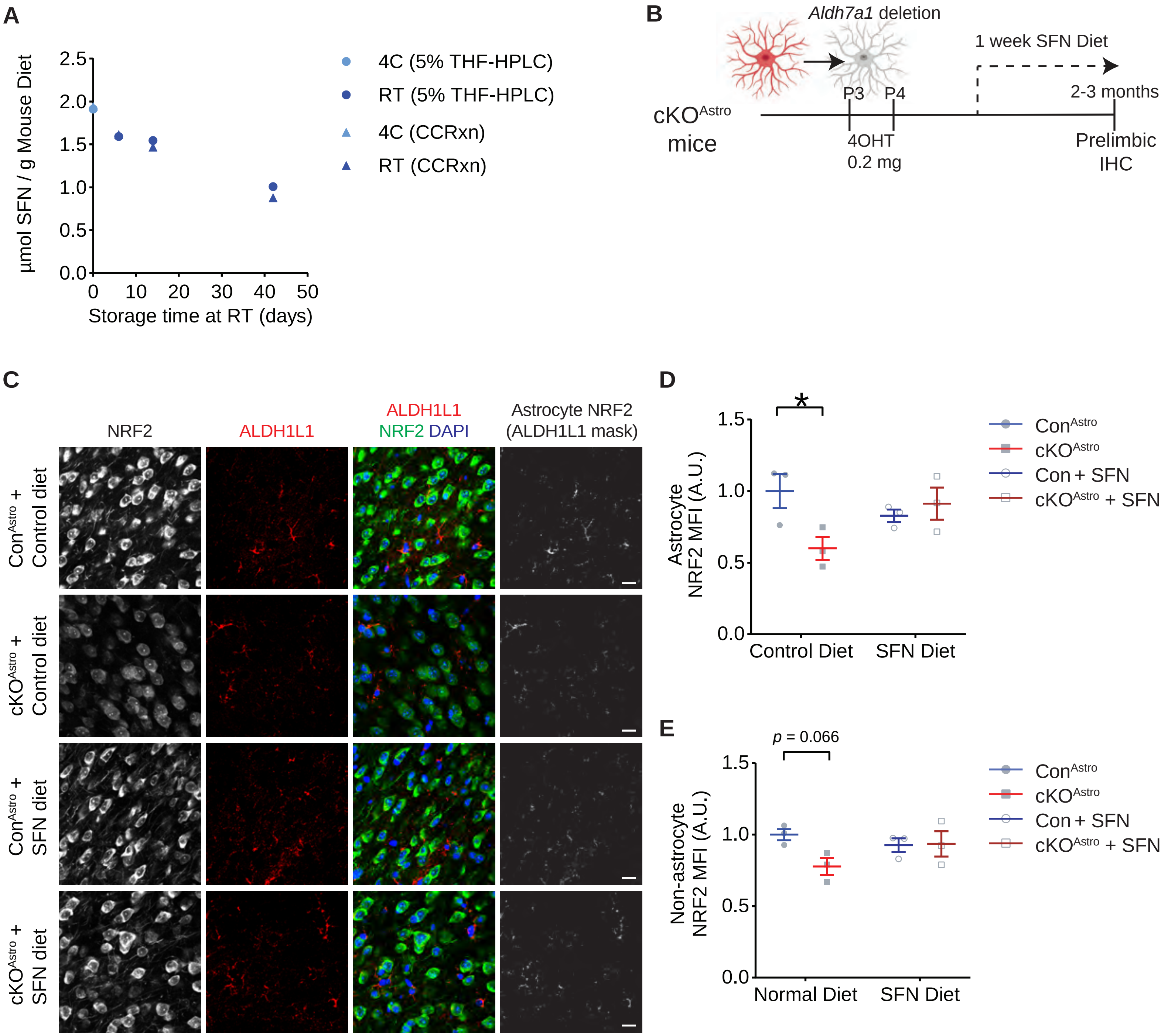

A
µmol SFN / g Mouse Diet
2.5
2.0
1.5
1.0
0.5
0.0
0
10
20
30
40
50
Storage time at RT (days)
4C (5% THF-HPLC)
RT (5% THF-HPLC)
4C (CCRxn)
RT (CCRxn)
B
Aldh7a1 deletion
1 week SFN Diet
cKOAstro mice
P3
P4
2-3 months
4OHT 0.2 mg
Prelimbic IHC
C
NRF2
ALDH1L1
ALDH1L1 NRF2 DAPI
Astrocyte NRF2 (ALDH1L1 mask)
ConAstro + Control diet
cKOAstro + Control diet
ConAstro + SFN diet
cKOAstro + SFN diet
D
*
Astrocyte NRF2 MFI (A.U.)
1.5
1.0
0.5
0.0
Control Diet
SFN Diet
ConAstro
cKOAstro
Con + SFN
cKOAstro + SFN
E
p = 0.066
Non-astrocyte NRF2 MFI (A.U.)
1.5
1.0
0.5
0.0
Normal Diet
SFN Diet
ConAstro
cKOAstro
Con + SFN
cKOAstro + SFN

**Fig. S4. Validation of sulforaphane diet.**

(A) Concentration of sulforaphane (SFN) in mouse diet pellets stored long-term at 4 °C and pellets left at room temperature for 6, 14, and 42 days. Individual data points measured by direct chromatography (%5 THF-HPLC) and by cyclocondensation (CCRxn) are shown.
(B) Schematic of 1-week dietary sulforaphane (SFN) treatment
(C-E) (C) Representative immunofluorescent images of NRF2 and astrocyte marker ALDH1L1 within the prelimbic cortex of $cKO^{Astro}$ and $Con^{Astro}$ mice on control diet or after 1 week on SFN diet. Right panels show NRF2 immunofluorescence in $ALDH1L1^{+}$ astrocytes (ALDH1L1 mask). Scale bars 20 µm. (D-E) Quantification of NRF2 mean fluorescent intensity (MFI) (D) in $ALDH1L1^{+}$ astrocytes (ALDH1L1 mask) and (E) outside $ALDH1L1^{+}$ astrocytes (inverse mask). 2-way repeated measures ANOVA with Holm-Sidak post-hoc tests: $n = 3$ mice; $*P < 0.05$, $P = 0.066$.
Data in (D-E) represent mean ± S.E.M.

**Fig. S5**

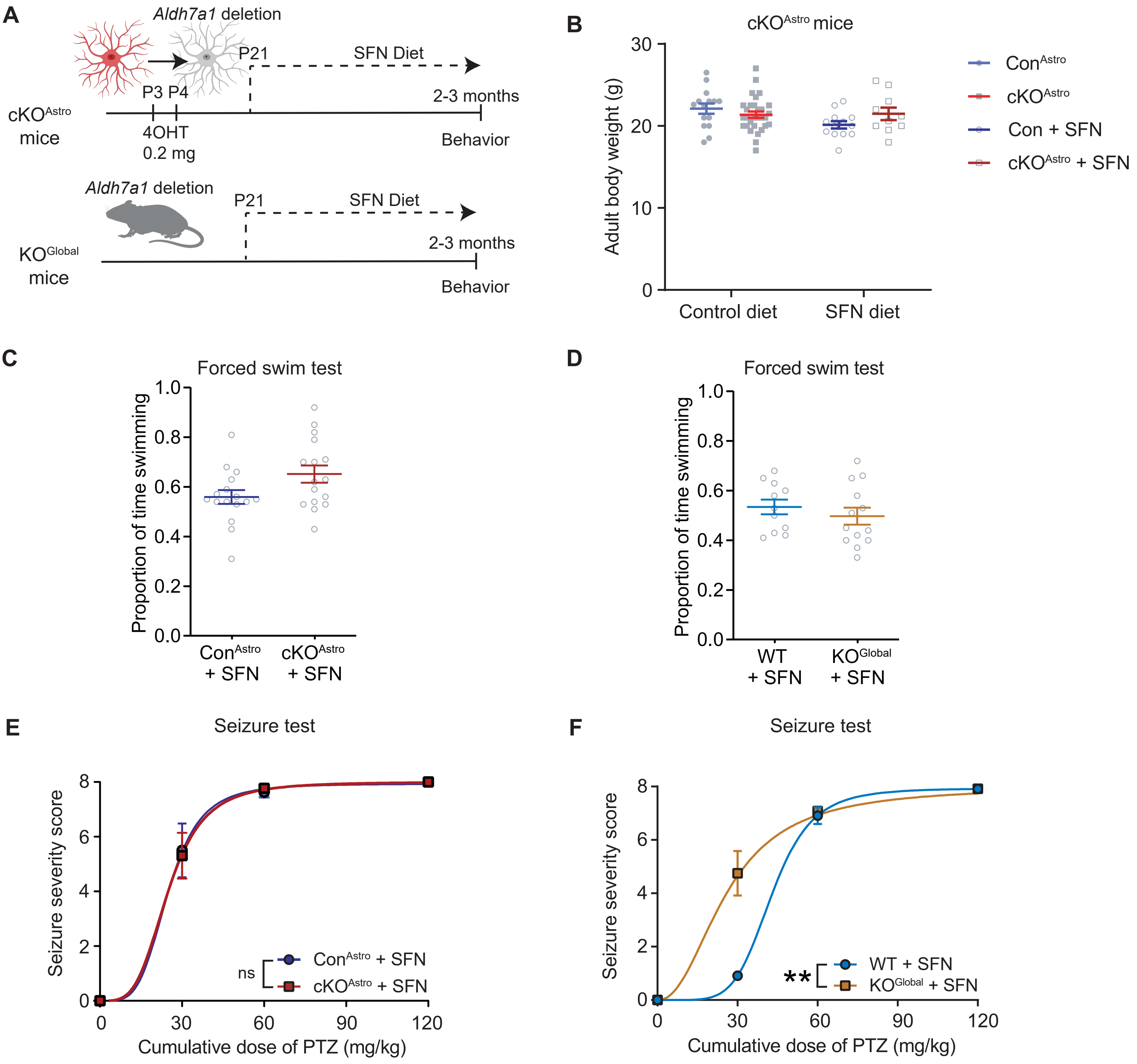

A
Aldh7a1 deletion
P21
SFN Diet
P3 P4
2-3 months
cKOAstro mice
4OHT 0.2 mg
Behavior
Aldh7a1 deletion
P21
SFN Diet
2-3 months
KOGlobal mice
Behavior
B
cKOAstro mice
Adult body weight (g)
30
20
10
0
Control diet
SFN diet
ConAstro
cKOAstro
Con + SFN
cKOAstro + SFN
C
Forced swim test
Proportion of time swimming
1.0
0.8
0.6
0.4
0.2
0.0
ConAstro + SFN
cKOAstro + SFN
D
Forced swim test
Proportion of time swimming
1.0
0.8
0.6
0.4
0.2
0.0
WT + SFN
KOGlobal + SFN
E
Seizure test
Seizure severity score
8
6
4
2
0
0
30
60
90
120
Cumulative dose of PTZ (mg/kg)
ns
ConAstro + SFN
cKOAstro + SFN
F
Seizure test
Seizure severity score
8
6
4
2
0
0
30
60
90
120
Cumulative dose of PTZ (mg/kg)
**
WT + SFN
KOGlobal + SFN

**Fig. S5. Behavioral analysis of sulforaphane diet.**

(A) Schematic of long-term dietary sulforaphane (SFN) treatment.
(B) Quantification of adult bodyweight in $cKO^{Astro}$ and $Con^{Astro}$ mice on control diet or long-term SFN diet (2-way ANOVA with Holm-Sidak post-hoc tests: $n = 10\text{-}30$ mice; $P > 0.05$).
(C-D) Quantification of the proportion of time swimming during the forced swim test by (C) $cKO^{Astro}$ and $Con^{Astro}$ mice on an SFN diet, (D) $KO^{Global}$ and WT mice on an SFN diet (Student's t-test: $n = 16$ mice ($cKO^{Astro}$), 11-13 mice ($KO^{Global}$); $P > 0.05$).
(E-F) Quantification of seizure severity in response to repeated doses of pentylenetetrazol (PTZ) during seizure threshold test in (E) $cKO^{Astro}$ and $Con^{Astro}$ mice on an SFN diet, (F) $KO^{Global}$ and WT mice on an SFN diet. Lines represent fitted dose-response curves. Extra sum-of-squares F test on dose-response $EC_{50}$: $n = 8\text{-}13$ mice ($cKO^{Astro}$), 11-12 mice ($KO^{Global}$); ns: $P > 0.05$, $^{**}P < 0.01$).
All data represent mean $\pm$ S.E.M.

**Fig. S6**

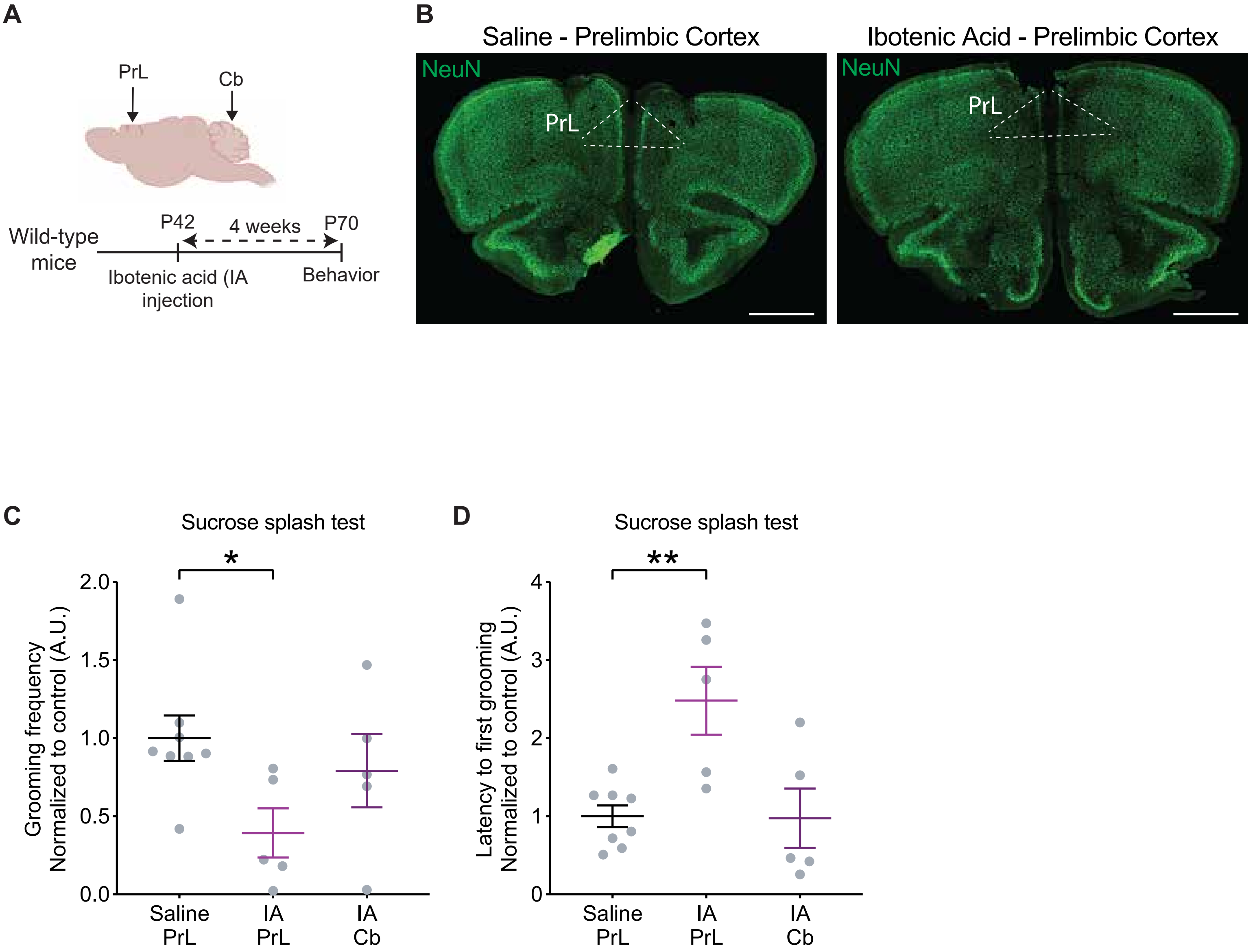
A
PrL
Cb
Wild-type mice
P42
4 weeks
P70
Ibotenic acid (IA injection
Behavior
B
Saline - Prelimbic Cortex
Ibotenic Acid - Prelimbic Cortex
NeuN
PrL
NeuN
PrL
C
Sucrose splash test
*
Grooming frequency Normalized to control (A.U.)
2.0
1.5
1.0
0.5
0.0
Saline PrL
IA PrL
IA Cb
D
Sucrose splash test
**
Latency to first grooming Normalized to control (A.U.)
4
3
2
1
0
Saline PrL
IA PrL
IA Cb

**Fig. S6. Pharmacological lesioning of prelimbic cortex impairs sucrose splash behavior.**

(A) Schematic of pharmacological lesioning with ibotenic acid.
(B) Representative images showing loss of NeuN immunolabeling in the prelimbic cortex of mice injected with ibotenic acid vs. saline-injected controls. Scale bars 1 mm.
(C-D) Quantification of (C) frequency of grooming events (D) and latency to first grooming during the sucrose splash test by mice injected in the prelimbic cortex (PrL) or cerebellum (Cb) with either saline or ibotenic acid (IA) by $KO^{Global}$ and WT mice (One-way ANOVA with Dunnett's post-hoc test: $n$ = 5-8 mice; $^{*}P < 0.05$, $^{**}P < 0.05$).
All data represent mean ± S.E.M.

### Fig. S7

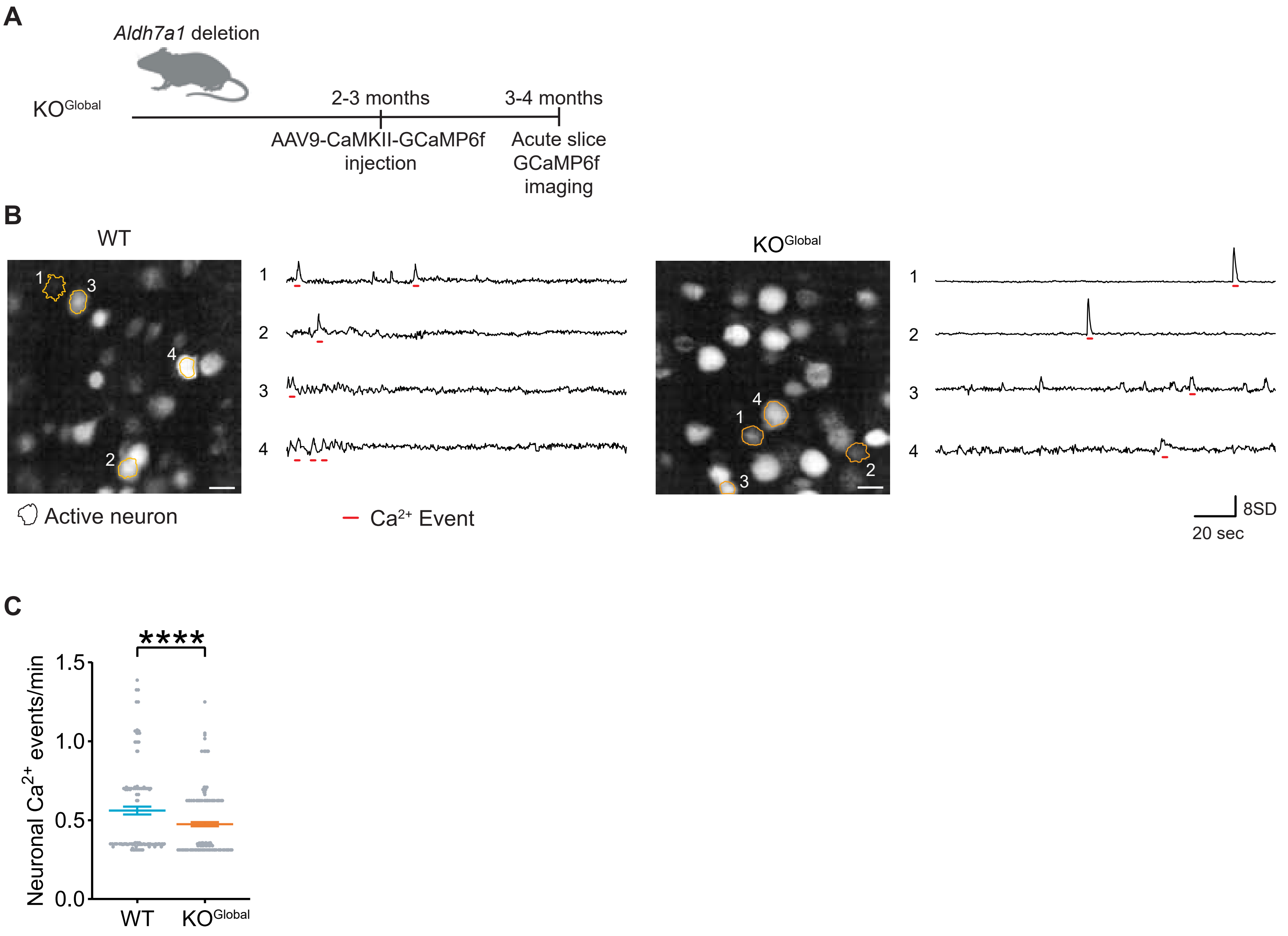
A
Aldh7a1 deletion
KOGlobal
2-3 months
3-4 months
AAV9-CaMKII-GCaMP6f injection
Acute slice GCaMP6f imaging
B
WT
KOGlobal
Active neuron
Ca2+ Event
8SD
20 sec
C
****
Neuronal Ca2+ events/min
1.5
1.0
0.5
0.0
WT
KOGlobal

**Fig. S7. Reduced layer 5 pyramidal neuron activity in the prelimbic cortex of $KO^{Global}$ mice.**

(A) Strategy to assess prelimbic layer 5 pyramidal neuron activity by GCaMP6f fluorescence in $KO^{Global}$ and WT mice.
(B) Representative images (left) and traces (right) of GCaMP6f fluorescence in the prelimbic cortex of $KO^{Global}$ and WT mice. Numbered orange outlines indicate neurons in images used for analysis and shown in traces. Events are indicated by red underlines. Scale bars, 20 μm.
(C) Quantification of prelimbic L5 pyramidal neuron $Ca^{2+}$ event frequency in WT and $KO^{Global}$ mice (Mann-Whitney test: $n$ = 139-209 neurons; ****$P < 0.0001$).
Data represent mean ± S.E.M.

Fig. S8

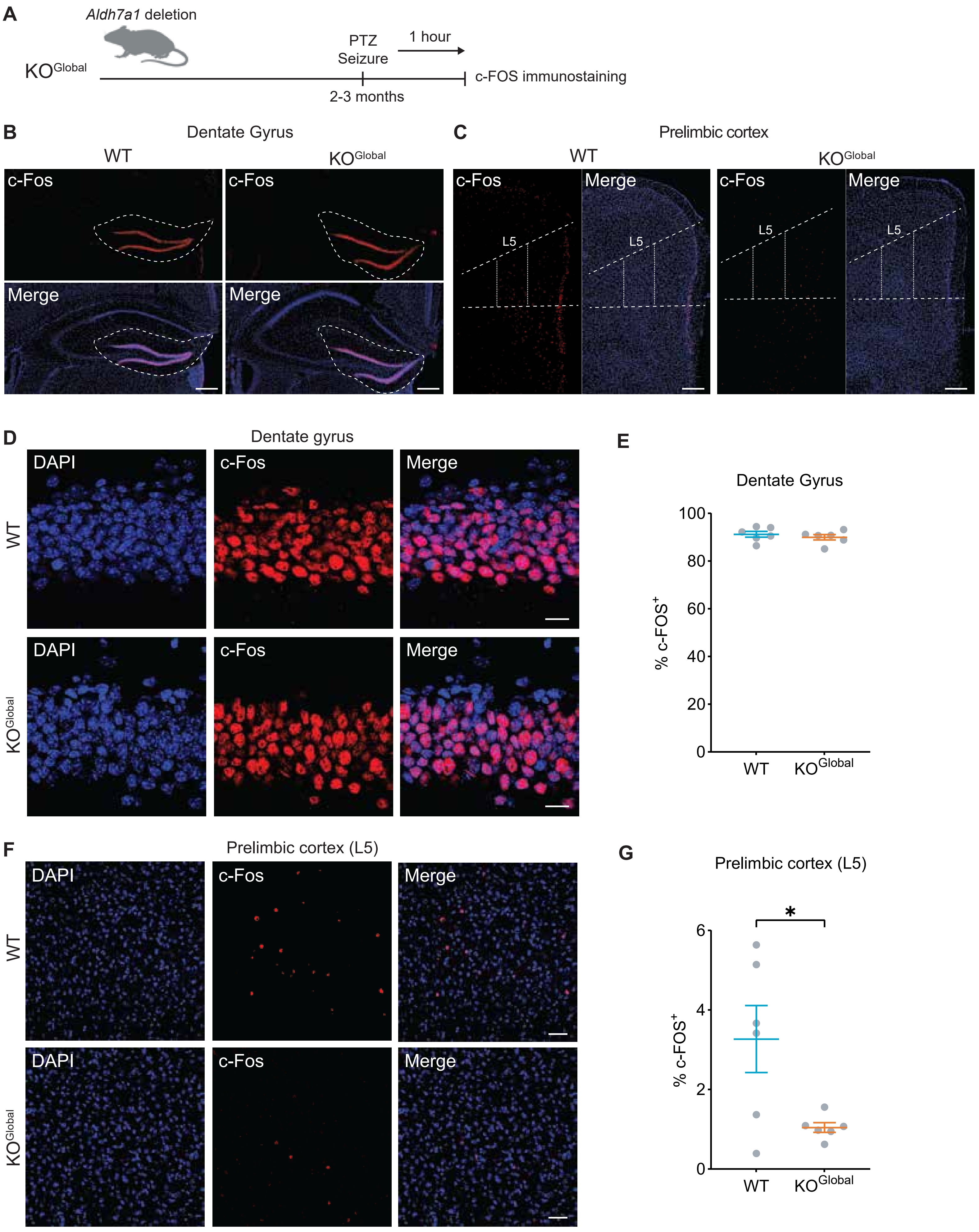

A
Aldh7a1 deletion
KOGlobal
PTZ
Seizure
1 hour
c-FOS immunostaining
2-3 months
B
Dentate Gyrus
WT
KOGlobal
c-Fos
Merge
C
Prelimbic cortex
L5
D
Dentate gyrus
DAPI
E
Dentate Gyrus
% c-FOS+
100
80
60
40
20
0
F
Prelimbic cortex (L5)
G
Prelimbic cortex (L5)
*
6
4
2

**Fig. S8. KO$^{Global}$ mice have a reduced percentage of c-FOS$^{+}$ cells in the prelimbic cortex during seizures.**

(A) Strategy to assess neuronal activity by c-FOS immunofluorescence in KO$^{Global}$ and wild-type (WT) mice at 1 hour after seizures induced by pentelyenetetrazol (PTZ).
(B-C) Representative images of c-FOS immunofluorescence in (B) the hippocampus and (C) the prelimbic cortex of KO$^{Global}$ and WT mice. White dashed outlines indicate the dentate gyrus in (B) and layer 5 (L5) prelimbic cortex in (C). Scale bars 200 μm.
(D-E) (D) Representative image and (E) quantification of c-FOS$^{+}$ cells in the dentate gyrus of KO$^{Global}$ and WT mice at 1 hour post-seizure (Student's t-test; n = 6 mice; $P > 0.05$).
(F-G) (F) Representative image and (G) quantification of c-FOS$^{+}$ cells in L5 prelimbic cortex of KO$^{Global}$ and WT mice at 1 hour post-seizure (Student's t-test; n = 6 mice; $^{*}P < 0.05$).
All data represent mean ± S.E.M.

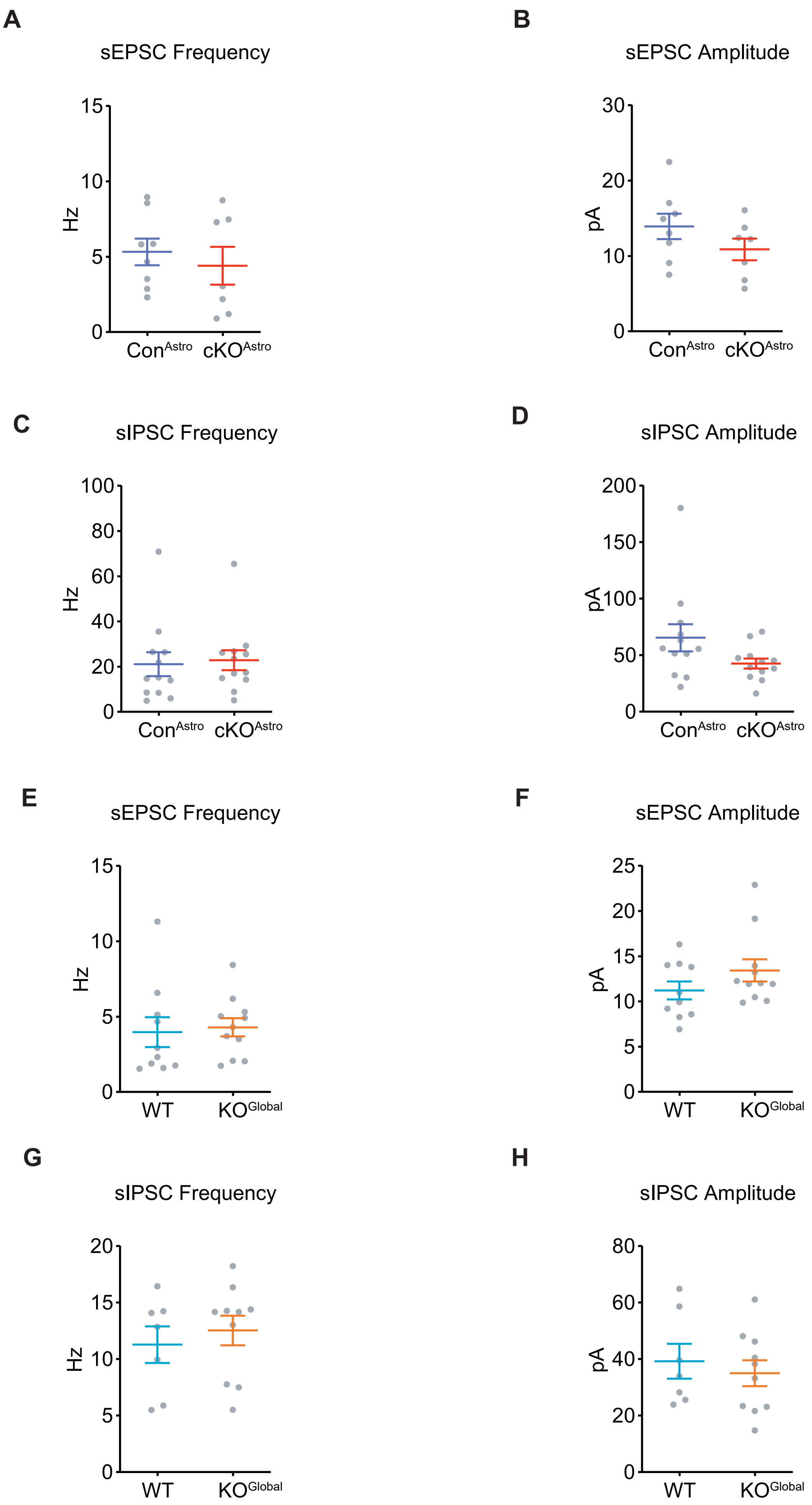
A
sEPSC Frequency
Hz
15
10
5
0
ConAstro
cKOAstro
B
sEPSC Amplitude
pA
30
20
10
0
ConAstro
cKOAstro
C
sIPSC Frequency
Hz
100
80
60
40
20
0
ConAstro
cKOAstro
D
sIPSC Amplitude
pA
200
150
100
50
0
ConAstro
cKOAstro
E
sEPSC Frequency
Hz
15
10
5
0
WT
KOGlobal
F
sEPSC Amplitude
pA
25
20
15
10
5
0
WT
KOGlobal
G
sIPSC Frequency
Hz
20
15
10
5
0
WT
KOGlobal
H
sIPSC Amplitude
pA
80
60
40
20
0
WT
KOGlobal

**Fig. S9. No changes in spontaneous postsynaptic currents on layer 5 pyramidal neurons in the prelimbic cortex of $cKO^{Astro}$ or $KO^{Global}$ mice.**

(A) Frequency of spontaneous excitatory postsynaptic currents (sEPSCs) in layer 5 pyramidal neurons in prelimbic cortex of $cKO^{Astro}$ and $Con^{Astro}$ mice (Student's t-test: $n$ = 7-8 cells; $P > 0.05$).
(B) Amplitude of sEPSCs in layer 5 pyramidal neurons in prelimbic cortex of $cKO^{Astro}$ and $Con^{Astro}$ mice (Student's t-test: $n$ = 7-8 cells; $P > 0.05$).
(C) Frequency of spontaneous inhibitory postsynaptic currents (sIPSCs) in layer 5 pyramidal neurons in prelimbic cortex of $cKO^{Astro}$ and $Con^{Astro}$ mice (Student's t-test: $n$ = 12 cells; $P > 0.05$).
(D) Amplitude of sIPSCs in layer 5 pyramidal neurons in prelimbic cortex of $cKO^{Astro}$ and $Con^{Astro}$ mice (Student's t-test: $n$ = 12 cells; $P > 0.05$).
(E) Frequency of sEPSCs in layer 5 pyramidal neurons in prelimbic cortex of $KO^{Global}$ and WT mice (Student's t-test: $n$ = 10-11 cells; $P > 0.05$).
(F) Amplitude of sEPSCs in layer 5 pyramidal neurons in prelimbic cortex of $KO^{Global}$ and WT mice (Student's t-test: $n$ = 10-11 cells; $P > 0.05$).
(G) Frequency of sIPSCs in layer 5 pyramidal neurons in prelimbic cortex of $KO^{Global}$ and WT mice (Student's t-test: $n$ = 7-10 cells; $P > 0.05$).
(H) Amplitude of sIPSCs in layer 5 pyramidal neurons in prelimbic cortex of $KO^{Global}$ and WT mice (Student's t-test: $n$ = 7-10 cells; $P > 0.05$).
All data represent mean ± S.E.M.

**Fig. S10**

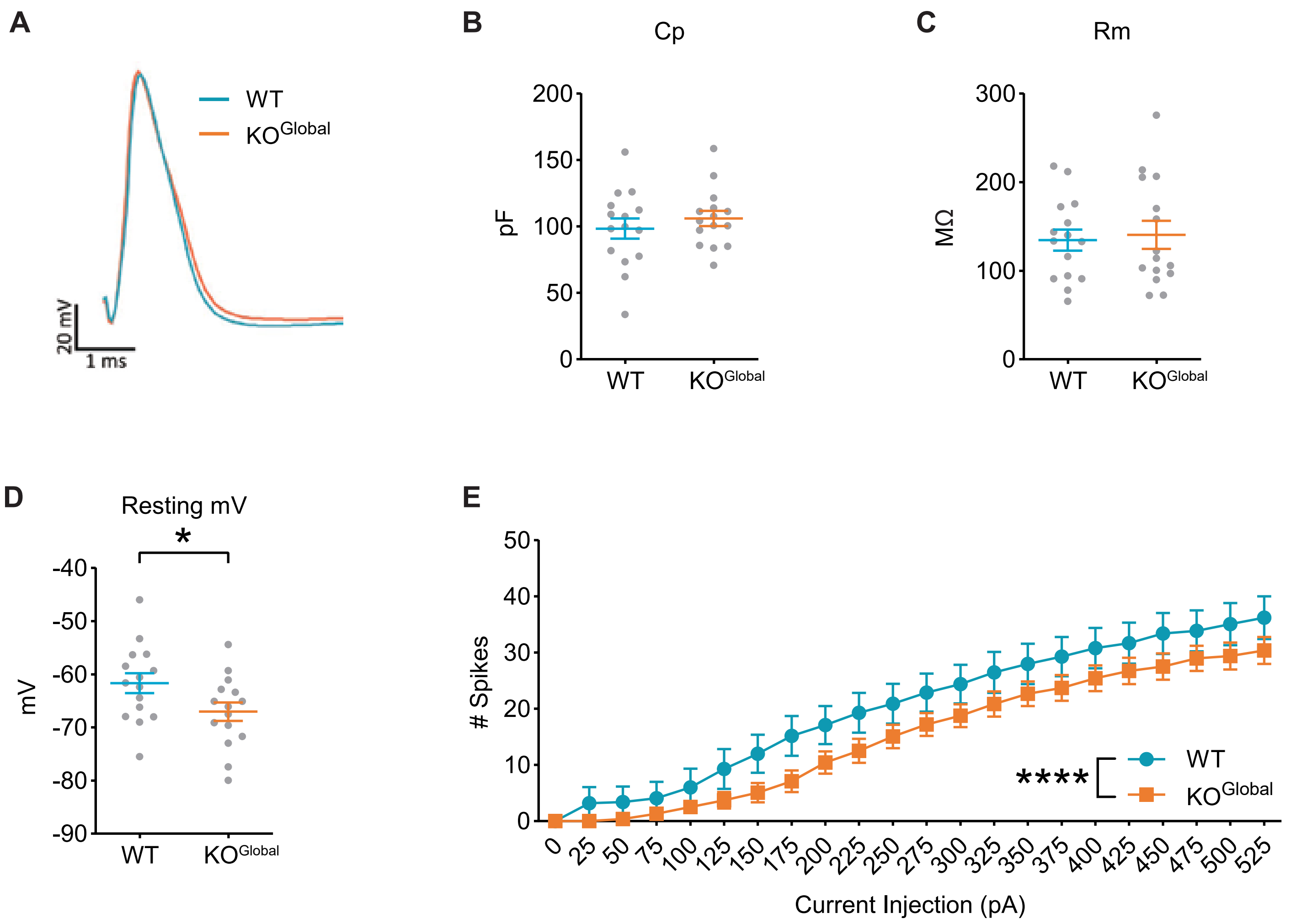
A
WT
KOGlobal
20 mV
1 ms
B
Cp
pF
200
150
100
50
0
WT
KOGlobal
C
Rm
MΩ
300
200
100
0
WT
KOGlobal
D
Resting mV
*
mV
-40
-50
-60
-70
-80
-90
WT
KOGlobal
E
### Spikes
50
40
30
20
10
0
****
WT
KOGlobal
0 25 50 75 100 125 150 175 200 225 250 275 300 325 350 375 400 425 450 475 500 525
Current Injection (pA)

**Fig. S10. Reduced resting membrane voltage and excitability of layer 5 pyramidal neurons in the prelimbic cortex of KO$^{Global}$ mice.**

(A) Representative action potential traces from prelimbic layer 5 pyramidal neurons in KO$^{Global}$ and WT mice.
(B-D) Quantification of (B) capacitance, (C) membrane resistance, and (D) resting membrane voltage of prelimbic layer 5 pyramidal neurons in KO$^{Global}$ and WT mice (Student's t-test: $n = 15$ cells; *$P < 0.05$).
(E) Quantification of the number of spikes generated per current injection in prelimbic layer 5 pyramidal neurons in KO$^{Global}$ and WT mice (2-way repeated measures ANOVA with Holm-Sidak post-hoc tests; $n = 10$-15 cells; main effect of genotype: **** $P < 0.0001$).
All data represent mean ± S.E.M.

### Fig. S11

**A**

ALDH7A1 Rescue Constructs

ALDH7A1 Rescue { AAV5-GfaABC1D-ALDH7A1-mCherry

Control AAV { AAV5-GfaABC1D-mCherry

Nrf2 Rescue Constructs

NRF2 Rescue { AAV5-GfaABC1D-Nrf2-mCherry + AAV5-GfaABC1D-mCherry-*Keap1* shRNA

Control AAVs { AAV5-GfaABC1D-mCherry + AAV5-GfaABC1D-mCherry-shRNA (scrambled)

**B**

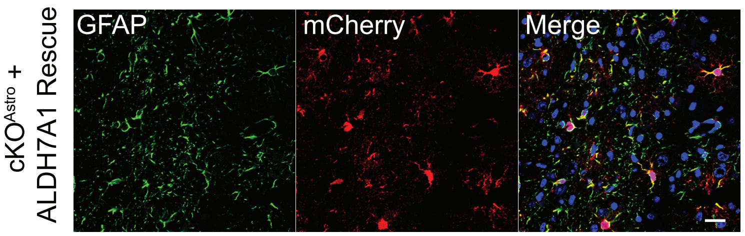


**C**

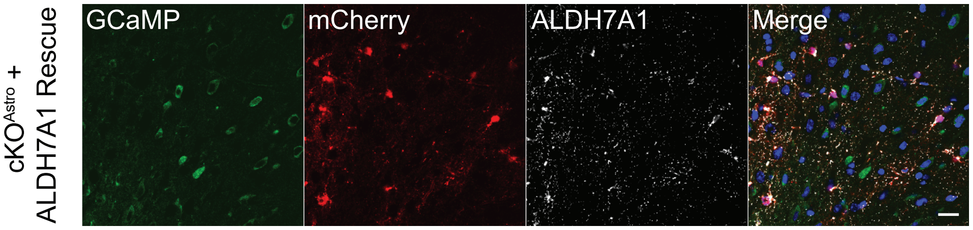


**D**

Electroporation of NRF2 Rescue plasmids

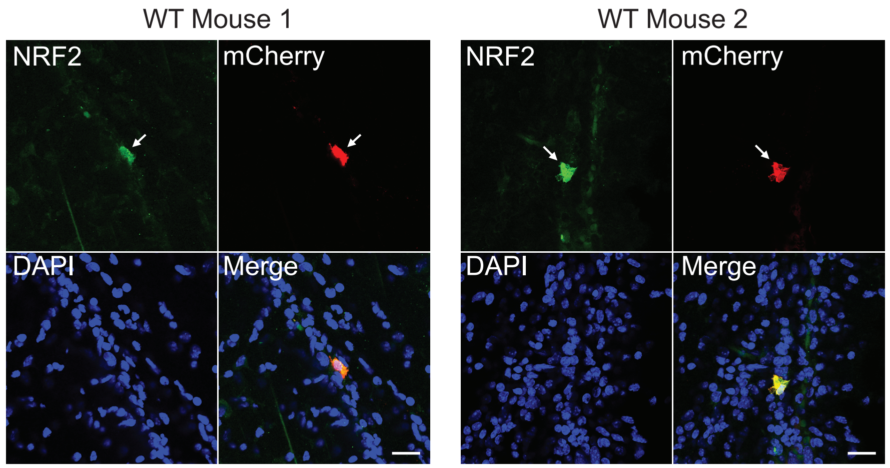

**Fig. S11. Validation of ALDH7A1 and NRF2 Rescue constructs.**

(A) Outline of ALDH7A1 and NRF2 Rescue constructs and respective controls.
(B-C) Representative images from the prelimbic cortex of $cKO^{Astro}$ mice injected with ALDH7A1 Rescue AAV (AAV5-GfaABC1D-ALDH7A1-mCherry) showing immunofluorescence of (B) astrocyte marker GFAP (green) and viral marker mCherry (red), and (C) GCaMP (green), ALDH7A1 (white), and viral marker mCherry (red). Scale bars 20 µm.
(D) Two representative images of NRF2 (green) and mCherry (red) immunofluorescence in wild-type (WT) mice electroporated with pZac2.1-GfaABC1D-Nrf2-mCherry and pZac2.1-GfaABC1D-mCherry-*Keap1* shRNA. Scale bars 20 µm.

**Fig. S12**

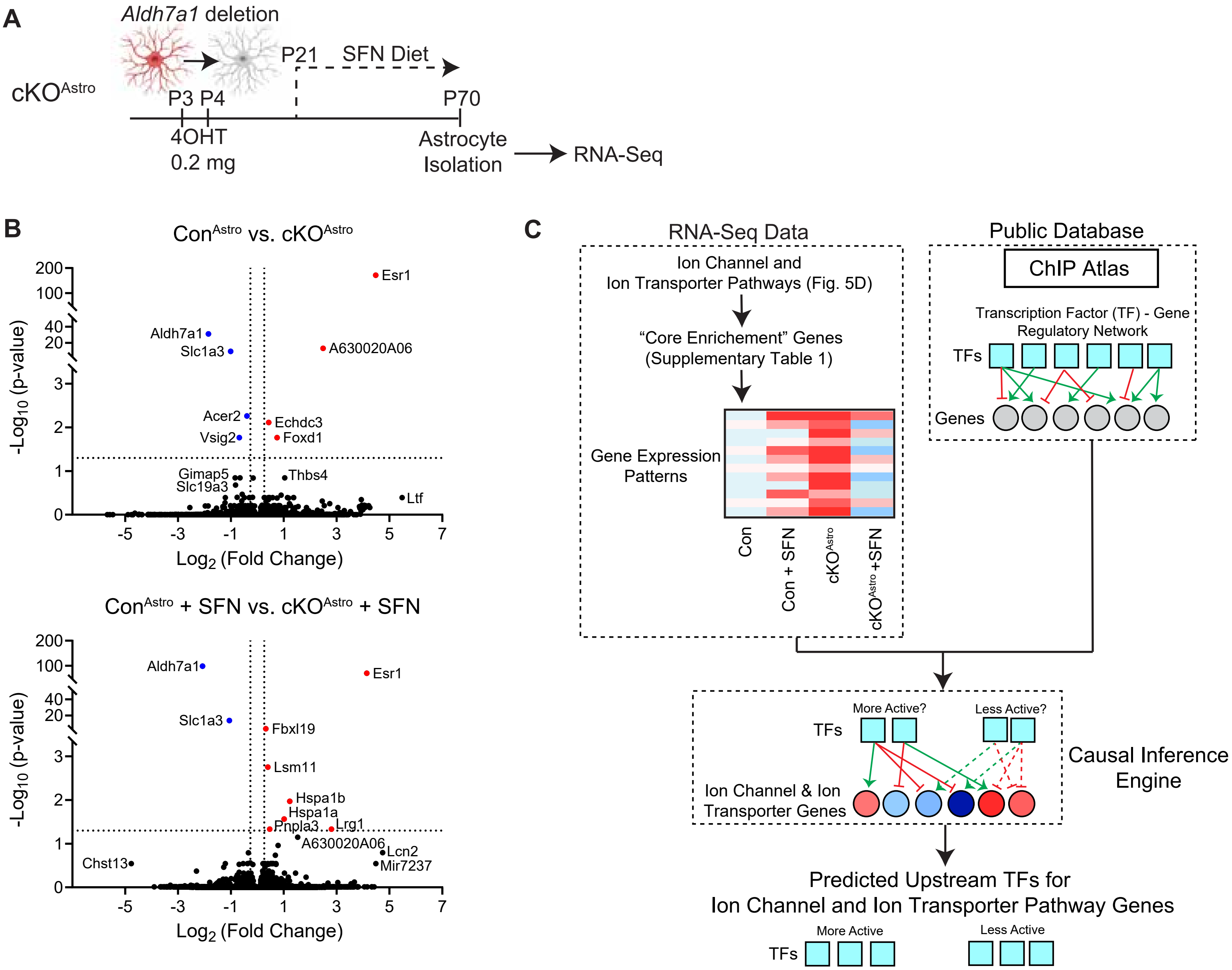


**D**

| Comparison | Predicted by CIE |
|---|---|
| cKO$^{Astro}$ vs. Con$^{Astro}$ | Yes |
| cKO$^{Astro}$ + SFN vs. Con$^{Astro}$ | No |
| Con$^{Astro}$ + SFN vs. Con$^{Astro}$ | No |

Predicted Upstream TFs for
Ion Channel and Ion Transporter Pathway Genes

More Active

ZBTB17, ZEB1, TET2, BRD2, GREB1, FOXF2, SMAD1

Less Active

GATA4

**Fig. S12. Putative upstream regulators of ion channel and ion transporter gene changes in cKO$^{Astro}$ astrocytes include redox-sensitive transcription factors.**

(A) Schematic of RNA-Sequencing of cortical astrocytes isolated from cKO$^{Astro}$, sulforaphane (SFN)-treated cKO$^{Astro}$ mice (cKO$^{Astro}$ + SFN), controls (Con$^{Astro}$), and SFN-treated controls (Con$^{Astro}$ + SFN).

(B) Volcano plots of RNA-Sequencing of astrocytes isolated from cKO$^{Astro}$ mice vs Con$^{Astro}$ mice and cKO$^{Astro}$ + SFN mice vs Con$^{Astro}$ + SFN. Dotted lines indicate cutoffs at $p < 0.05$ and $\log_2$ fold change > | 1.2 |.

(C) Visual summary of causal inference engine (CIE) analysis. As inputs, CIE used the gene expression patterns of "core enrichment" ion channel and ion transporter genes in astrocytes obtained via RNA-Seq as well as the ChIP Atlas public database of transcription factor (TF) – Gene regulatory networks. Based on these two inputs, CIE was used to make predictions about changes in the activity of TFs upstream of ion channel and ion transporter genes in each condition.

(D) Table indicates the strategy used to identify TFs affected in cKO$^{Astro}$ astrocytes and rescued by SFN. TFs fitting these criteria are listed below.
